# Viral Infection Induces Alzheimer’s Disease-Related Pathways and Senescence in iPSC-Derived Neuronal Models

**DOI:** 10.1101/2025.06.11.659008

**Authors:** Hana Hribkova, Veronika Pospisilova, Katerina Amruz Cerna, Tereza Vanova, Jan Haviernik, Jiri Sedmik, Ondrej Bernatik, Jaroslav Olha, Jan Raska, Sona Cesnarikova, Miriam Satkova, Klara Plesingrova, Petra Strakova, Andrea Fortova, Katarina Gresova, Manolis Maragkakis, Michaela Sadibolova, Rudolf Kupcik, Ivo Fabrik, Marie Vajrychova, Katerina Sheardova, Daniel Ruzek, Dasa Bohaciakova

**Affiliations:** Department of Histology and Embryology, Faculty of Medicine, Masaryk University, Kamenice 5, Brno 625 00, Czech Republic; International Clinical Research Center, St. Anne’s University Hospital Brno, Pekarska 53, Brno 602 00, Czech Republic; Veterinary Research Institute, Hudcova 296/70, Brno 621 00, Czech Republic; Department of Experimental Biology, Faculty of Science, Masaryk University, Kamenice 5, Brno 625 00, Czech Republic; Institute of Parasitology, Biology Centre of the Czech Academy of Sciences, Branisovska 1160/31, Ceske Budejovice 370 05, Czech Republic; Laboratory of Genetics and Genomics, National Institute on Aging, Intramural Research Program, National Institutes of Health, Baltimore, MD 21224, USA; Biomedical Research Centre, University Hospital Hradec Kralove, Sokolska 581, Hradec Kralove 500 05, Czech Republic

**Author notes:** Corresponding author: Dasa Bohaciakova, Department of Histology and Embryology, Faculty of Medicine, Masaryk University, Kamenice 5, Brno 625 00, Czech Republic. School of Biological Sciences, University of Canterbury, Christchurch 8041, New Zealand (Jan Haviernik). Authors contributed equally (Hana Hribkova, Veronika Pospisilova, Katerina Amruz Cerna, Tereza Vanova).

**Keywords:** Cerebral organoids, Herpes virus, Tick-borne Encephalitis, Alzheimer’s disease, Senescence

## Abstract

**INTRODUCTION:** The Pathogen Infection Hypothesis proposes that β-Amyloid (Aβ) functions as an antimicrobial peptide, with pathogen-induced aggregation potentially contributing to Alzheimer’s disease (AD) pathology.

**METHODS:** We used human iPSC-derived 2D neurons and 3D cerebral organoids from wild-type and familial AD (*PSEN1/2* mutant) lines to model acute infections with HSV-1 and TBEV and Aβ aggregation. Transcriptomic and proteomic analyses were conducted to assess molecular responses.

**RESULTS:** HSV-1, but not TBEV, induced robust Aβ clustering, which was, however, dependent on extracellular amyloid peptides. Transcriptomic profiling revealed widespread HSV-1-induced changes, including activation of neurodegeneration-related pathways. Proteomic profiling confirmed enrichment of neurodegeneration- and senescence-associated secretome signatures. *PSEN1/2* mutations did not alter the acute infection response. Reanalysis of independent datasets confirmed our findings and revealed a limited protective effect of acyclovir.

**DISCUSSION:** Results directly support the Pathogen Infection Hypothesis and suggest that preventing viral infections via vaccinations may represent a feasible approach to reducing AD risk.

## 1. Background

Alzheimer’s disease (AD) is a progressive neurodegenerative disorder and the most common cause of dementia worldwide. It is pathologically characterized by the accumulation of extracellular β-Amyloid (Aβ) plaques, intracellular neurofibrillary tangles composed of hyperphosphorylated TAU, neuroinflammation, synaptic dysfunction, and eventual neuronal loss [1,2]. Despite decades of intensive research, the precise triggers for Aβ deposition and the cascade of molecular events that lead to neurodegeneration remain incompletely understood. Although most AD cases are sporadic, familial forms caused by *APP, PSEN1, PSEN2,* or *SORL1* gene mutations underscore the importance of amyloid precursor protein (APP) processing and Aβ production in disease pathogenesis [3–5].

The Pathogen Infection Hypothesis has recently gained renewed attention [6,7]. This hypothesis proposes that Aβ peptides may act as evolutionarily conserved antimicrobial peptides that are produced and aggregated in response to pathogen invasion as part of the brain’s innate immune defense [8]. In this model, Aβ entraps and neutralizes microbes, such as herpes simplex virus type 1 (*Simplexvirus human alpha 1;* HSV-1), through rapid aggregation. This initially protective mechanism may become maladaptive when chronic or excessive, leading to plaque formation and sustained inflammation [9,10]. Supporting this view, several studies have detected HSV-1 DNA in post-mortem brains of AD patients, particularly in regions affected by the disease [11–13], and herpes zoster vaccination has recently been associated with reduced AD risk in population-based studies [14]. Importantly, growing evidence from population studies further suggests that repeated antiviral vaccinations may have protective effects against AD and related health outcomes. Specifically, influenza vaccination in older adults has been associated with a reduced risk of developing AD, with the strength of this effect increasing with the number of repeated vaccinations [15]. Similarly, a higher number of COVID-19 vaccine doses has been linked to better self-reported health outcomes, including improved mental health, in individuals experiencing long COVID [16]. Nevertheless, the experimental evidence remains mixed, and critical questions persist. Not all studies have replicated the findings of HSV-1 involvement in AD [17,18], and other neurotropic viruses, especially those causing acute infections of the neural system, have not been systematically evaluated for their potential to induce similar responses. Additionally, the mechanisms by which pathogens influence AD-related molecular pathways, such as protein aggregation, oxidative stress, endoplasmic reticulum (ER) stress, and cellular senescence, are not fully elucidated. Whether these responses are modulated by the genetic background of the host, particularly familial AD mutations, is also poorly understood.

To address these gaps, recent efforts have focused on human-relevant *in vitro* models to study early molecular events in the human context, such as induced pluripotent stem cell (iPSC)-derived neurons and 3D cerebral organoids (COs) [19–21]. These models allow the recapitulation of aspects of brain development, neural differentiation, and disease- associated phenotypes in a controlled environment, free from the limitations of post- mortem tissue or non-human models. In this study, we leveraged iPSC-derived 2D neuronal cultures and 3D COs derived from both wild-type (WT) individuals and from patients carrying *PSEN1* or *PSEN2* mutations to systematically examine the cellular and molecular responses to two distinct neurotropic viruses: i) HSV-1, causing chronic infections, and ii) tick-borne encephalitis virus (*Orthoflavivirus encephalitidis*; TBEV), causing acute infections. We aimed to investigate whether these viruses induce features of AD pathology, such as Aβ aggregation, neuroinflammatory responses, and senescence, and whether these responses differ depending on viral type or host genotype. Using transcriptomic, proteomic, and imaging analyses, we explored the relevance of infection-driven mechanisms in the context of neurodegeneration and AD.

## 2. Methods

### 2.1 Cell Culture and Differentiation

In this study, we used previously established and well-characterized human iPSC lines, including both WT and familial AD patient-derived lines (*PSEN1/2* mutant) [22,23], as detailed in **Table A**. All iPSCs were cultured in mTeSR medium (85850, StemCell Technologies) on Matrigel-coated plates (734-1440, Corning) and routinely passaged using TrypLE (12605036, Thermo Fisher Scientific) or 0.5 mM EDTA (AM9260G, Thermo Fisher Scientific). These lines were subsequently differentiated into inducible 2D neurons and 3D COs, as described below. The exact number of organoids and cell lines used in each experimental replicate is provided in **Table B**.

#### 2.1.1 Inducible Neuronal Differentiation (2D Cultures)

Rapid differentiation of 2D neurons was performed using NGN2 transgenic WT iPSCs (RRID:CVCL_C7XJ), following a previously described protocol [23]. Doxycycline (D9891, Merck) was used to induce NGN2 expression during the first three days. On day 3, cells were replated at a density of 0.166 × 10⁶ cells/cm² onto coverslips coated with poly-L-ornithine (P3655, Merck) and Laminin (23017015, Gibco), where they were allowed to mature before viral infection experiments.

#### 2.1.2 Cerebral Organoid Differentiation (3D Cultures)

To generate COs, both WT and *PSEN1/2* mutant iPSC lines [22] were differentiated using a modified version of the Lancaster *et al.* protocol [19], as previously optimized by our group [24]. Organoids were used for viral infection studies at days 53 and 93 of development (with embryoid body seeding designated as day 0). For experiments with pre-treatment of amyloid peptides, organoids were cultured in a medium supplemented with 100 nM synthetic Aβ40 (A1075, Merck) or Aβ42 (A9180, Merck) peptides for seven days (from day 46 to day 53) prior to viral exposure.

### 2.2 Viral Infection

Two viruses (HSV-1 and TBEV) were used to infect mature neurons and COs. HSV-1 viral particles (strain MacIntyre) were produced using Vero cells (kidney epithelial cells from African green monkeys), as previously described [25]. The virus was kindly provided by Prof. Andreas Sauerbrei, German Reference Laboratory for HSV and VZV, Germany. TBEV, including a construct expressing a stable mCherry reporter, as well as the TBEV strain Hypr (used as a control strain without mCherry reporter in selected experiments with 2D neurons), was generated using porcine kidney stable (PS) cells, following a previously established protocol [26]. The Hypr strain was provided by the Collection of Arboviruses, Biology Centre of the Czech Academy of Sciences (https://arboviruscollection.bcco.cz).

#### 2.2.1 Viral Infection of Inducible 2D Neurons

On day 17 of differentiation (D17), NGN2-induced neurons were exposed to HSV-1 (multiplicity of infection, MOI 0.0001), mCherry-TBEV or TBEV Hypr at multiple MOIs (0.0001, 0.01, 0.1, 1) for 30 minutes. Viruses were diluted in viral infection medium composed of DMEM (LM-D1112, Biosera), 10% fetal bovine serum (FB-1001, Biosera), 1% penicillin-streptomycin (XC-A4110, Biosera), and 1% L-glutamine (XC-T1755, Biosera). After infection, cells were washed and maintained in the fresh neuronal medium for four additional days. Control neurons were treated with the viral medium without viral particles.

#### 2.2.2 Viral Infection of Cerebral Organoids

COs were infected either on day 53 (D53, early timepoint) or day 93 (D93, late timepoint) of differentiation. HSV-1 (MOI 0.0001) or mCherry-TBEV (MOI 0.1), diluted in the viral infection medium, was applied for 24 hours. After the infection, organoids were transferred to fresh CO differentiation medium (COII) and cultured for up to 7 days post-infection (dpi). Control organoids underwent the same procedure but were not exposed to the virus.

### 2.3 Immunochemistry and Microscopic Analysis

Immunocytochemistry and imaging were conducted on both 2D neurons and 3D CO sections. Inducible neurons were fixed and labeled as previously described [27] with antibodies against Aβ and infection-specific and nuclear markers, as detailed in **Table C**. Samples were visualized using a Zeiss Axio Imager.Z1 widefield microscope, and tiled images were captured to quantify Aβ accumulation. For COs, fixed tissue was embedded in agarose, sectioned at 250 μm thickness using a vibratome, and processed for immunohistochemistry. Organoid sections were cleared using glycerol/fructose to improve imaging depth and quality.

Microscopy of cleared organoid sections was performed using a Zeiss Axio Observer.Z1 microscope with LSM 800 confocal unit. Fluorescent signals from labeled proteins and nuclei were acquired using high-resolution confocal settings and appropriate laser/filter combinations. Aβ plaques were identified and quantified using Imaris software (v9.8.2), and volumetric data were analyzed using GraphPad Prism (v9). Full protocols and antibody lists are available in the **Supplementary Methods** and **Table C**.

### 2.4 Real-time quantitative PCR, Western Blot, and ELISA

RNA, protein, and secreted peptide analyses were used to characterize infection-related molecular changes in COs. For viral gene detection using real-time quantitative PCR (qRT-PCR), COs were UV-inactivated and processed for total RNA extraction using a Direct-zol RNA Microprep Kit (R2062, ZymoResearch). RNA was reverse-transcribed to cDNA and analyzed by qRT-PCR (LightCycler 480, Roche) using virus-specific primers verified previously [20,28] and listed in **Table D**. Ct values were calculated automatically, and no viral gene expression was detected in non-infected controls.

For protein-level analysis using Western blotting, COs were lysed and processed using a standard protocol [29]. Protein concentrations were measured using the DC Protein Assay (5000112, Bio-Rad), separated by SDS-PAGE, transferred to PVDF membranes, and probed with specific antibodies (**Table C**). Signal was detected using Clarity Max Western ECL Substrate (1705062, BioRad) and imaged with a ChemiDoc system.

Finally, to quantify secreted Aβ40 and Aβ42 peptides using ELISA, cell cultures of 2D neurons at D21 and COs at D57 and D97 were incubated in Essential 6 Medium for an additional 3 days. The media were analyzed using commercial ELISA kits (KHB3544, KHB3482, Thermo Fisher Scientific), with normalization based on total protein content measured from corresponding total cell lysates to account for COs size variability. All measurements were performed in biological and technical replicates (see **Table B**). Complete protocols and antibody/primer lists are available in the **Supplementary Methods** and **Table C**.

### 2.5 RNA Isolation, 3’mRNA-sequencing and Data Analysis

For transcriptomic analysis, total RNA was extracted from single organoids using the Direct-zol RNA Microprep Kit (R2062, ZymoResearch) after UV inactivation and PBS washing (see **Supplementary Methods** for full protocol). Biological replicates from six independent iPSC lines (WT 1, 2, 3; *PSEN1/2* mutants 1, 2, 3) were pooled per condition (non-treated=NTR, HSV-1, TBEV). Only RNA samples with RINe≥ 7.5 were processed for library preparation using the QuantSeq FWD 3’mRNA Library Prep Kit (Lexogen), and sequencing was performed on a NextSeq 500 system (Illumina), generating ∼10 million reads per sample.

Sequencing data were processed in R (v4.4.2) [30]. Raw reads were trimmed, quality-checked, and aligned to the GRCh38 human genome [31] using STAR v2.7.0 [32]. Gene counts were generated via htseq-count [33] and analyzed with DESeq2 v1.46.0 [34] to identify differentially expressed genes (DEGs) in between infected and corresponding non-infected controls (p.adj<0.1, log2FC>|0.5|, baseMean>100). Downstream visualization included volcano plots and principal component analysis, with top DEGs selected by weighted score [log2FC*-log10(p.adj)]. Functional enrichment analysis was performed using clusterProfiler v4.14.3 [35,36] and visualized with ggplot2 v3.5.1 [37], UpSetR v1.4.0 [38] and enrichplot v1.26.2 [39]. Targeted analyses were based on preselected GOBP terms connected to neurodegeneration and HSV-1 infection [40–43] (**Table E**) using msigdbr package v7.5.1 [44]. Further methodological detail is provided in the **Supplementary Methods**.

### 2.6 Protein Isolation, Mass Spectrometry and Data Analysis

Proteomic processing of conditioned media from COs followed a rigorous pipeline detailed in the **Supplementary Methods**. Proteins from the Essential 6 culture medium harvested from COs infected with HSV-1, TBEV and non-infected controls (media from 146 individual COs) were precipitated, quantified, and subjected to enzymatic digestion using a combination of lysyl endopeptidase and trypsin. Peptides were labeled with Tandem Mass Tag (TMT) 16plex reagents and pooled into multiplexes that included global internal standards (GIS) to control batch variability. Designed TMT multiplexes were subsequently fractionated under high-pH conditions and analyzed by liquid chromatography-tandem mass spectrometry (LC-MS/MS) using Q Exactive Plus instrument.

MS/MS spectra were processed in Proteome Discoverer 3.0 and proteins were quantified across 146 secretomes. Three samples were excluded from further analysis due to low protein content. TMT intensities were normalized within and across multiplexes to correct artificial technical variability. Differentially secreted proteins (DSPs) were identified using LIMMA with Benjamini-Hochberg correction (FDR<0.05), and proteins with p.adj<0.1 and |log2FC|>0.5 were selected for further analysis. These DSPs were mapped to ENTREZ IDs and functionally characterized using over-representation analysis as described in the **Supplementary Methods** (section 5.1). Full methodological details are available in **Supplementary Methods**.

### 2.7 Data visualization and availability

Schematic representations in this study were created in BioRender. Data from qRT-PCR, Western blot, ELISA and microscopic analysis were visualized using GraphPad Prism (version 9.0 for Windows, GraphPad Software, www.graphpad.com). The 3’mRNA-seq data were deposited in NCBI’s Gene Expression Omnibus [45] and are accessible through GEO Series accession number GSE295890 and link https://www.ncbi.nlm.nih.gov/geo/query/acc.cgi?acc=GSE295890. The proteomic secretome datasets are available in the ProteomeXchange Consortium via the PRIDE partner repository [46] with the dataset identifier PXD063530 and link https://www.proteomexchange.org/.

## 3. Results

### 3.1 Differential Accumulation of Amyloid Beta in 2D Neuronal Cultures Following HSV-1 and TBEV Infections

Previous research has indicated that various viral and microbial pathogens, including HSV-1, could lead to Aβ accumulation [47–49]. Indeed, protein aggregation as a mechanism of antimicrobial protection represents one of the cornerstones of the Pathogen Infection Hypothesis [6]. However, the effect of TBEV infection, despite its neurotropism, remained unclear. Thus, during initial experiments, we assessed whether viral infections by both HSV-1 and TBEV promote the formation of Aβ clusters in our 2D human iPSC-derived neuronal cultures. We utilized WT, karyotypically normal iPSCs, differentiated into TUJ and MAP2-positive neurons with mature morphology via NGN2 overexpression [23] (**Figure 1A**). As schematized in **Figure 1B**, after 17 days of culture, we infected the neurons with live HSV-1 or TBEV at a MOI 0.0001 for 30 minutes, as previously reported for HSV-1 [50,51]. Following this incubation period, excess viruses were washed out, and cultivation continued for four days with media changes every two days. Non-infected neurons of the same age, undergoing the same experimental handling and media changes, were used as control. Infection was assessed using specific viral markers (i.e., ICP4 for HSV-1 and mCherry reporter for TBEV; **Figure 1C**) and validated by Western blotting and qRT-PCR (**Figure A.1**). Notably, HSV-1 infection led to a significant increase in Aβ signal in the 2D cultured iPSC-derived neurons, as previously described [52,53]. In contrast, TBEV infection did not induce a widespread increase in the Aβ signal compared to NTR controls, even when different MOIs or TBEV strains were tested (**Figure A.2**).

**Figure 1:**
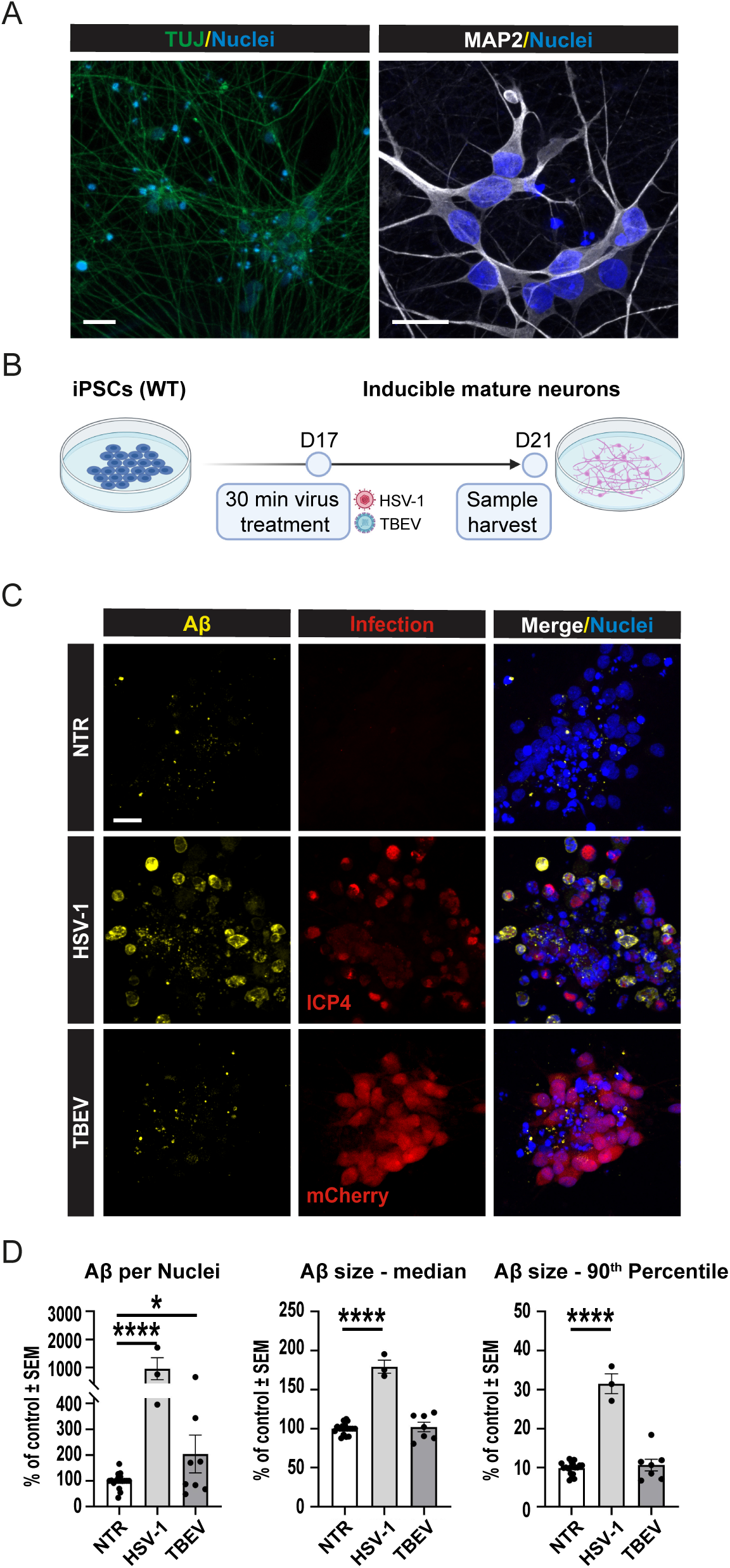
Differential accumulation of Aβ in 2D neuronal cultures following HSV-1 and TBEV infections. **(A)** WT, karyotypically normal iPSCs, differentiated into TUJ-positive and MAP2-positive neurons with mature morphology via NGN2 overexpression. Scale bars = 20μm. (**B)** Scheme of the experiment. After 17 days of culture, neurons were infected with live HSV-1 or TBEV at a MOI 0.0001 for 30 minutes. Following this incubation period, the virus was washed out, and cultivation continued until D21 with media changes every two days. **(C)** Representative pictures of NTR neurons or infected with HSV-1 or TBEV. Infection by HSV-1 (ICP-4) and TBEV (mCherry reporter) was detected using Immunofluorescence staining (red) along with detection of Aβ (yellow), and nuclei (blue) and **(D-F)** quantified. We assessed **(D)** volume of Aβ particles normalized to the volume of cell nuclei per image; **(E)** median volume of Aβ; and **(F)** 90^th^ percentile of Aβ particle size. Each dot represents one biological replicate, n≥3, significance was evaluated using unpaired t-test, error bars represent mean ±SEM; *p<0.05, ****p<0.001; scale bar = 20 µm. See **Table B** for reference on specific number of samples, replicates and cell line details.

Image analysis quantification (**Figures 1D-F**) further supported these findings. We specifically quantified i) the overall quantity of Aβ signal per cell (assessed by the volume of Aβ particles normalized to the volume of cell nuclei per image; **Figure 1D**), ii) changes in the overall size of all Aβ particles (visualized as the median volume of Aβ; **Figure 1E**) and iii) specifically the number of large Aβ clusters (quantified as the 90^th^ percentile of Aβ particle size; **Figure 1F**). Our analyses confirmed that all of these parameters were increased in neurons infected with HSV-1 compared to the controls, including the quantity of Aβ per cell, the median size of all Aβ particles, and the amount of large Aβ clusters. These changes were highly significant (****p<0.001), confirming previous findings by other groups [47,52,53], as well as validating our 2D neuronal model. In contrast, TBEV infection had a significantly milder effect on the neuronal cultures, not affecting the formation of large Aβ clusters or the median size of Aβ, although we did detect an increased volume of Aβ signal per cell nuclei upon TBEV infection (*p<0.05). We conclude that, compared to HSV-1 infection, TBEV did not induce the same widespread response that would lead to significant Aβ clustering in 2D neuronal cultures.

### 3.2 Acute Pathogen Infection of 3D Cerebral Organoids Does Not Result in Accumulation of Aβ

Prompted by initial findings from 2D neuronal cultures, we investigated how 3D COs derived from iPSCs responded to viral infections by HSV-1 and TBEV. Importantly, previous studies that used neural stem cell- or neuron-based 3D models reported APP accumulation following HSV-1 infection [47,51]. However, two independent studies that used self-organizing iPSC-derived COs reported conflicting results [52,54], with one study showing the accumulation of Aβ after HSV-1 infection, whereas the other study did not detect Aβ clusters in 3D COs. Moreover, the response of 3D organoids to TBEV infection has not yet been explored.

Thus, to address these questions, we employed our previously established and characterized protocol for generating COs from iPSCs and subsequent image analysis and evaluation of Aβ signal from histological sections [24]. As shown schematically in **Figure 2A**, we differentiated three independent WT iPSC lines into COs. Upon reaching maturity (D53), the organoids were infected with either HSV-1 or TBEV. After 24 hours, COs were washed and cultured for an additional 6 days (until D60), with media changes every 2 days. To examine the impact of the aging of COs and the associated natural accumulation of Aβ on the response of organoids to viral infection, we repeated the same experimental setup using D93 organoids, harvesting them at D100.

**Figure 2:**
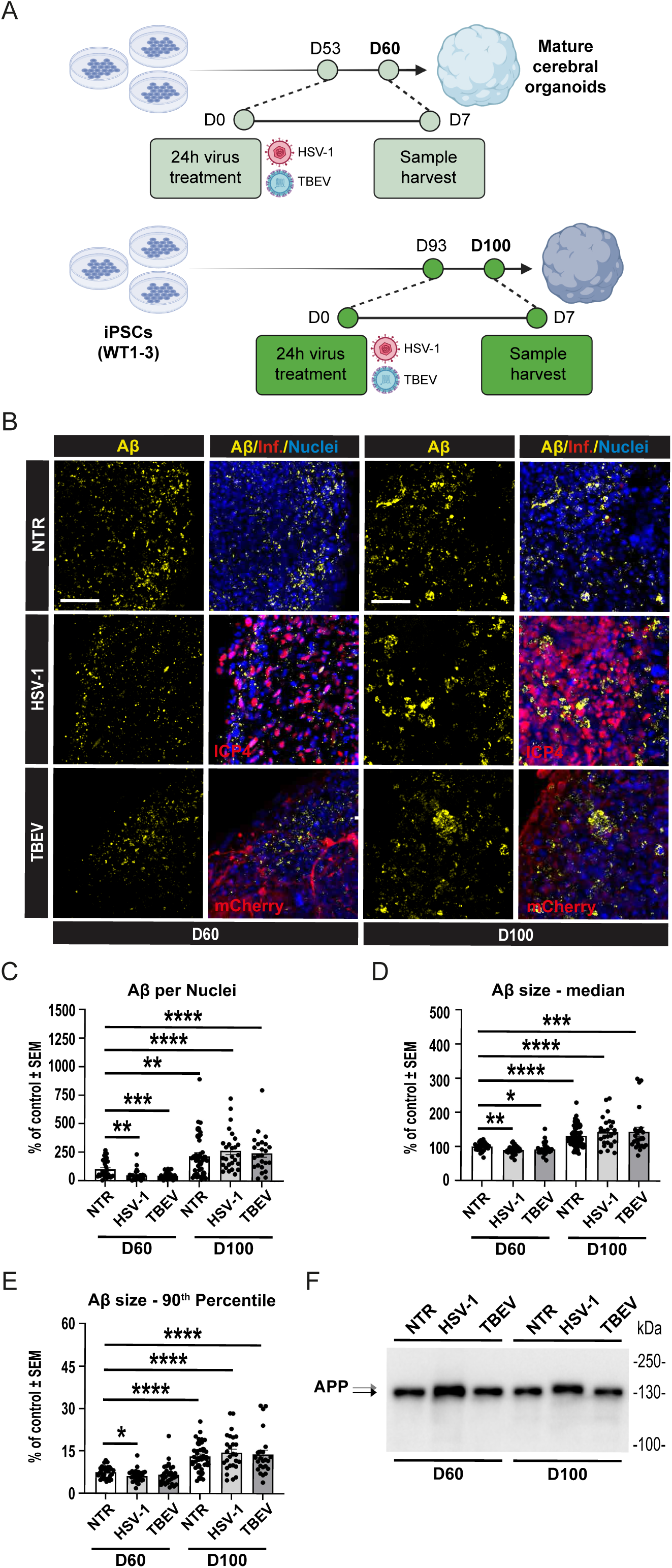
Acute pathogen infection of 3D COs does not result in Aβ accumulation. **(A)** Scheme of experimental workflow. Three independent WT iPSC lines were differentiated into COs. On day D53, COs were infected with either HSV-1 or TBEV. After 24 hours, organoids were washed and maintained in culture until D60, with medium changes every two days. A parallel experimental setup was performed with older organoids infected at D93 and harvested at D100. **(B)** Representative immunofluorescence images showing Aβ staining in COs at D60 and D100 after infection with HSV-1 or TBEV. Scale bars = 50 µm. **(C–E)** Image analysis and quantification of Aβ signal after infection. We assessed **C)** volume of Aβ particles normalized to the volume of cell nuclei per image; **(D)** median volume of Aβ; and **(E)** 90^th^ percentile of Aβ particle size. Each dot represents one biological replicate, n≥3, significance was evaluated using unpaired t-test, error bars represent mean ±SEM; ****p<0.001, **p < 0.01. See **Table B** for reference on specific number of samples, replicates and cell line details. **(F)** Representative Western blot analysis of APP. A consistent band shift in APP full-length protein was observed following HSV-1 infection, but not TBEV infection, in both D60 and D100 COs.

The results of qRT-PCR analysis confirmed successful infection of organoids by both HSV-1 and TBEV compared to NTR controls (**Figure B**). However, unlike in 2D neuronal cultures, viral infection of our 3D COs did not result in increased Aβ signal or clustering (**Figure 2B**). As illustrated in **Figure 2B** and quantified from a total of 172 CO sections in **Figures 2C–E**, infection of D60 organoids did not increase the volume of Aβ signal per cell nuclei, the median size of Aβ, or the formation of large Aβ clusters. Importantly, D100 COs displayed significantly elevated Aβ parameters compared to D60 controls, confirming our earlier observations [24]. Nevertheless, despite this maturation-associated increase in Aβ, viral infection with either HSV-1 or TBEV did not alter Aβ parameters compared to NTR controls. This indicates that even advanced organoid maturation, accompanied by a significant increase in all measured Aβ parameters within organoids, does not promote Aβ clustering in response to infection by either HSV-1 or TBEV.

It is of note that while detecting full-length APP in infected and non-infected CO samples using Western blotting, we observed a small but consistent shift in the APP band following HSV-1 infection but not TBEV infection (**Figure 2F**). Previous studies have shown that HSV-1 can induce specific cleavage or phosphorylation of APP fragments in mouse and human models [55,56]. However, to our knowledge, this is the first report suggesting a potential post-translational modification of full-length endogenous APP in response to HSV-1 infection. Importantly, it suggests that although APP clustering is not induced by HSV-1, the HSV-1 infection indeed alters the intracellular biology of APP, although its nature and consequences remain to be explored.

### 3.3 With Abundant Aβ Peptides in the Extracellular Space, 3D Cerebral Organoids Respond to HSV-1 Infection, But Not TBEV Infections, by Forming Aβ Clusters

During our initial experiments with 2D and 3D neuronal models, we observed markedly different responses to viral infections. This discrepancy led us to investigate why stem cell-derived 2D and 3D neuronal models yielded such divergent outcomes, particularly given that previous studies using 3D systems frequently reported Aβ clustering following HSV-1 infection [47,51,53]. Notably, those studies were primarily based on 3D models utilizing differentiating neural stem cells or mature neurons embedded in scaffolds, matrices, or cultured as spheroids. In contrast, two studies employing self-organizing iPSC-derived COs reported conflicting results [52,54]. These observations prompted us to examine the levels of extracellular (secreted) Aβ peptides in our culture systems, which may influence Aβ seeding and the formation of APP clusters after HSV-1 infection. Interestingly, our analyses revealed that COs secrete substantially lower amounts of Aβ peptides into the culture medium compared to 2D neuronal cultures. As shown in **Figure C**, standard 2D neuronal cultures secreted, on average, 5.585 pg of Aβ40 and 0.150 pg of Aβ42 per 1 µg of total protein, whereas standard cultures of COs released only 0.463 pg of Aβ40 and 0.011 pg of Aβ42 per 1 µg of protein at D60 and 0.615 pg of Aβ40 and 0.008 pg of Aβ42 per 1 µg of protein at D100. These data indicate approximately 10-fold lower basal levels of extracellular Aβ peptides in COs and suggest that the availability of Aβ in the extracellular space is a critical factor for Aβ seeding and its proposed antimicrobial activity, independent of whether a 2D or 3D culture system is used.

Thus, to test the hypothesis that elevated extracellular Aβ peptides can induce APP cluster formation in a 3D organoid system, we supplemented CO cultures with exogenous Aβ peptides. Specifically, as schematized in **Figure 3A**, three independent WT iPSC lines were differentiated into COs. Beginning at D46, synthetic Aβ40 or Aβ42 peptides were added to the culture medium for seven days. At D53, the Aβ-enriched medium was washed out, and the organoids were infected with HSV-1 or TBEV. After 24 hours, COs were washed again and maintained in culture for an additional six days, with media changes every two days. Importantly, as shown in **Figure 3B**, COs pre-treated with soluble Aβ40 or Aβ42 exhibited a widespread increase in Aβ signal in CO sections following HSV-1 infection. This effect was not observed after TBEV infection or in non-infected samples. Quantitative analysis of microscopy data quantified from a total of 207 CO sections (**Figures 3C–E**) confirmed that Aβ pre-treatment combined with HSV-1 infection significantly increased the number of large Aβ clusters (**p<0.01). Additionally, Aβ42, but not Aβ40, pre-treatment led to a significant increase in Aβ signal per cell and in median cluster size (*p<0.05). Consistent with previous observations in 2D neuronal cultures shown in **Figure 1**, these effects were not detected after TBEV infection or in organoids treated with Aβ peptides alone. These results thus support the hypothesis that an excess of extracellular Aβ peptides is sufficient to induce Aβ cluster formation upon HSV-1 infection, regardless of the used culture system. Furthermore, they suggest that this process largely relies on the availability of extracellular Aβ peptides capable of seeding Aβ aggregation.

**Figure 3:**
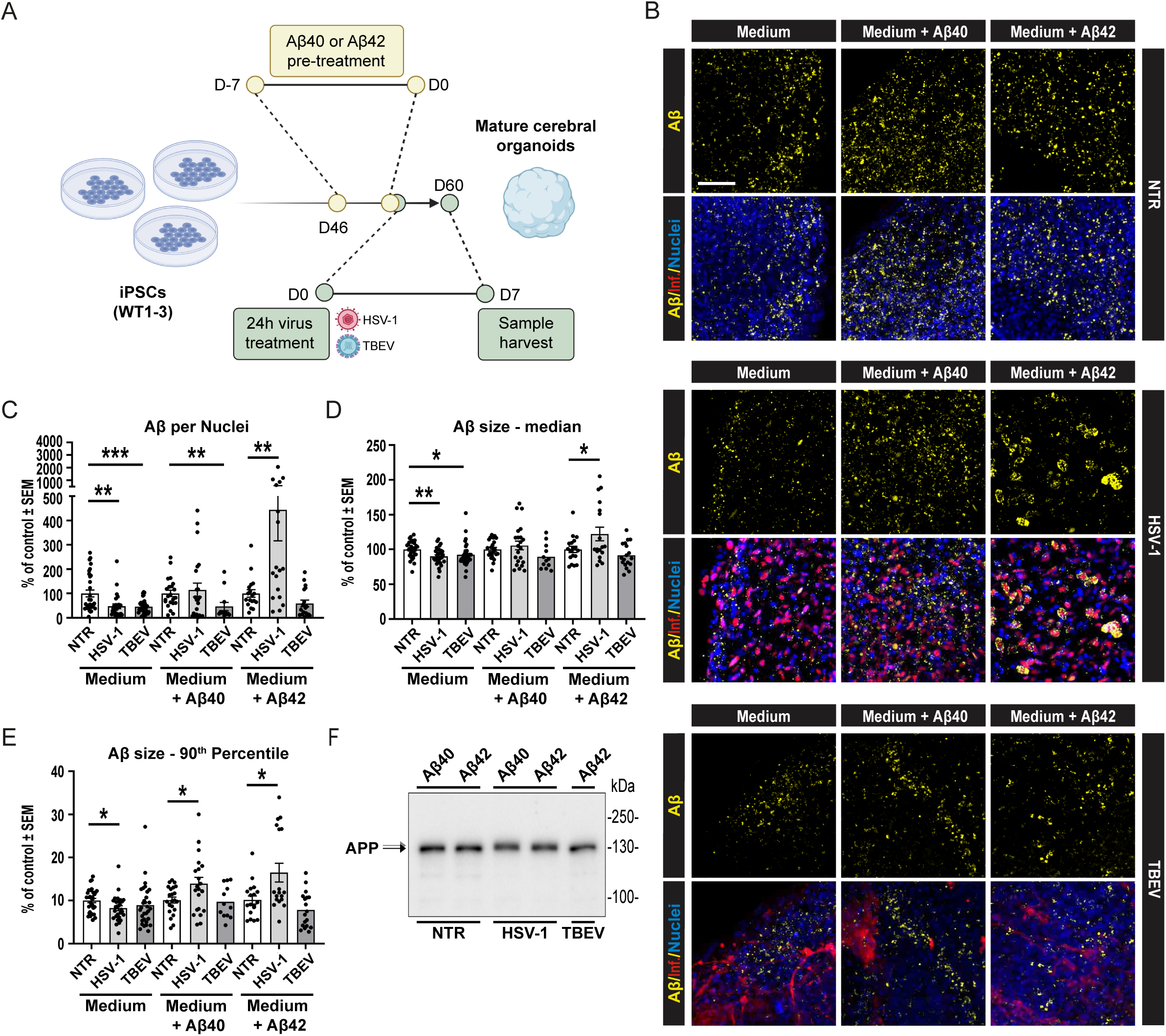
Exogenous Aβ peptide supplementation enables Aβ cluster formation in COs following HSV-1, but not TBEV, infection. **(A)** Schematic overview of the experimental setup. Three independent WT iPSC lines were differentiated into COs. From D46, organoids were treated with synthetic Aβ40 or Aβ42 peptides for seven days. At D53, Aβ peptides were washed out, and organoids were infected with HSV-1 or TBEV. After 24 hours, COs were washed and cultured in the fresh medium until D60. **(B)** Representative immunofluorescence images showing increased Aβ signal in COs pre-treated with Aβ peptides and infected with HSV-1 and TBEV. Scale bar = 50μm. **(C–E)** Image analysis and quantification of Aβ signal after infection. We assessed **(C)** volume of Aβ particles normalized to the volume of cell nuclei per image; **(D)** median volume of Aβ; and **(E)** 90^th^ percentile of Aβ particle size. Each dot represents one biological replicate, n≥3, significance was evaluated using unpaired t-test, error bars represent mean ±SEM; ***p<0.005, **p<0.01, *p<0.05. **(F)** Western blot analysis of full-length APP. See **Table B** for reference on specific number of samples, replicates and cell line details.

Notably, and irrespectively of Aβ cluster formation or Aβ40/Aβ42 pre-treatment, Western blot analysis of full-length APP again revealed a small but consistent band shift (**Figure 3E**), similar to that shown in **Figure 2F**. This shift was observed exclusively following HSV-1 infection, indicating that HSV-1, but not TBEV, may influence the post-translational modification of APP, independently of extracellular Aβ-mediated seeding.

### 3.4 Transcriptomic Profiling Reveals Distinct Molecular Responses to HSV-1 and TBEV Infection in Cerebral Organoids

Our results thus far indicated that the formation of Aβ clusters in organoids occurs exclusively in the presence of abundant amyloid peptides. These, functioning as antimicrobial agents, target HSV-1 viral particles [48], a response not observed with the TBEV virus. However, the Pathogen Infection Hypothesis encompasses both 1) a protein aggregation as a mechanism of antimicrobial protection and 2) a molecular response, which mainly involves the inflammatory response that may cause neuronal damage over time, but that also causes direct damage and cell death [6]. What molecular changes underlie the reaction of organoids to viral infection in our experimental setup remained unaddressed.

To investigate this molecular response using our organoid model, we performed 3’mRNA sequencing (3’ mRNA-seq) to examine gene expression changes in organoids harvested at D60. The analysis included three independent WT iPSC-derived organoid lines harvested 7 dpi with either HSV-1 or TBEV and corresponding non-infected controls, following the infection scheme depicted in **Figure 2A**. Four individual organoids per cell line (WT1, 2, 3) per condition (NTR, HSV-1, TBEV) were combined into nine pooled samples and analyzed. Principal component analysis revealed that HSV-1 treatment significantly altered the global gene expression profile, leading to extensive changes, whereas TBEV infection caused more modest shifts (**Figure 4A**). Differential expression analysis as illustrated in volcano plots confirmed that HSV-1 treatment resulted in the substantial deregulation of over 1 600 genes (p.adj<0.1, log2FC>|0.5|, see **Supplementary Methods**), with most of these genes being upregulated compared to non-infected controls (**Figure 4B, top**). Interestingly, subsequent unbiased ORA using the KEGG database identified that the genes impacted by HSV-1 infection are involved in multiple pathways linked to neurodegeneration, including AD, Parkinson’s disease, amyotrophic lateral sclerosis, and prion diseases (**Figure 4C**). The top 20 upregulated and downregulated genes (see **Supplementary Methods**) within these categories are depicted in a heatmap (**Figure D**) and contribute to the regulation of key cellular processes implicated in AD pathology. Notably, several upregulated genes (e.g., *PIK3C3, ATF6, APAF1, TARDBP*/*TDP-43*) are involved in stress responses, protein aggregation, or programmed cell death, while downregulated genes (e.g., *NDUF* genes*, UBA1, PSMD4*) reflect mitochondrial dysfunction and impaired proteasomal clearance, hallmarks of early neurodegenerative changes. All deregulated genes are listed in **Table F**. In contrast, TBEV infection only significantly affected 22 genes (p.adj<0.1, log2FC>|0.5|, see **Supplementary Methods**, **Figure 4B, bottom**), upregulating genes solely associated with the antiviral immune response (**Figure 4C**). These included key elements of the interferon signaling pathway, such as *STAT1*, a crucial transcription factor, and *MX1*, an interferon-induced GTPase that defends against a broad array of viruses. Additional upregulation was observed in numerous interferon-induced downstream genes, such as *IFIT* and *IFI* genes, which are vital for antiviral defense.

**Figure 4:**
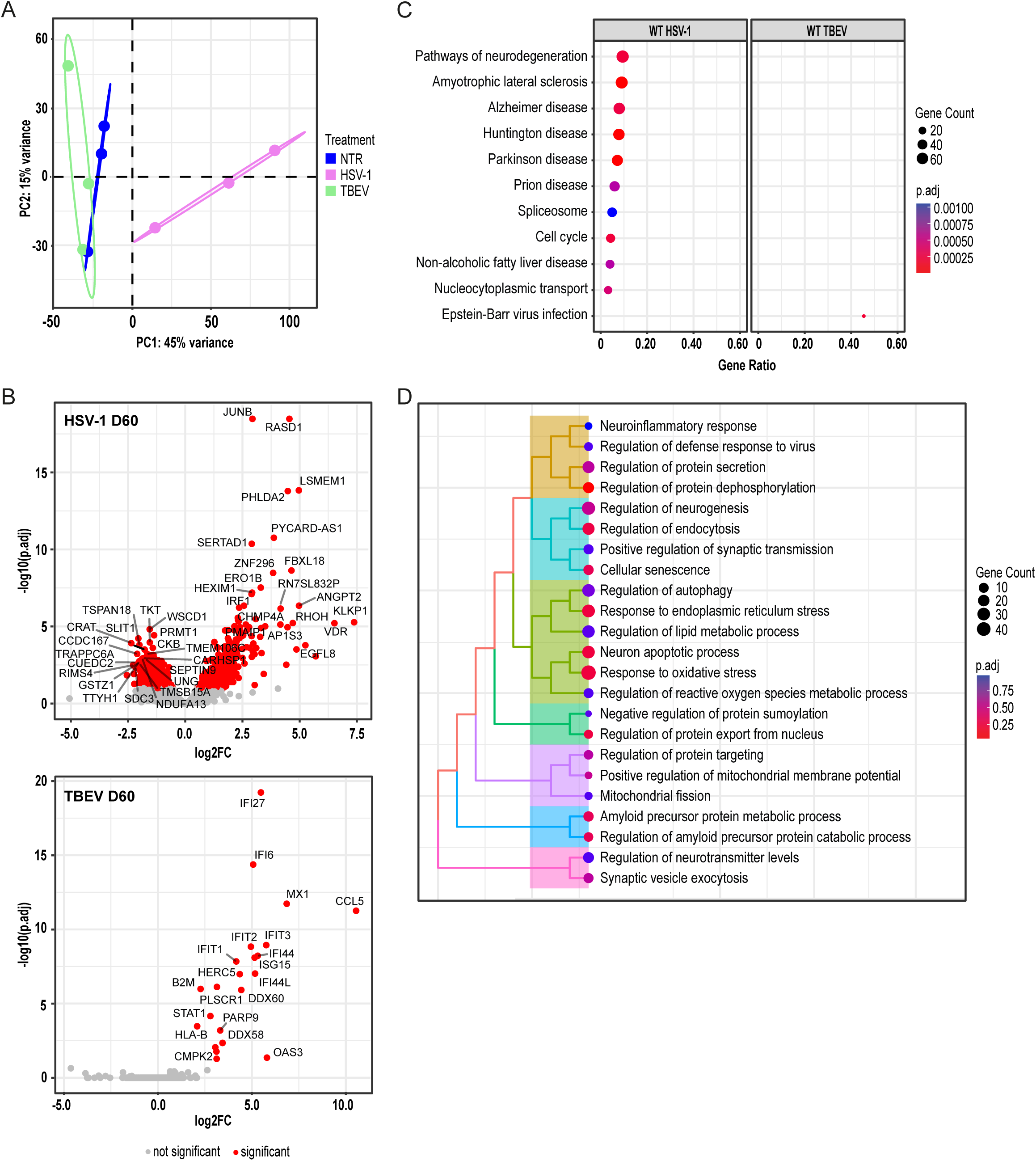
Transcriptomic profiling reveals distinct molecular responses to HSV-1 and TBEV infection in COs. **(A)** Principal component analysis of transcriptomic data from WT iPSC-derived COs harvested at D60 following 7-day infection with HSV-1 or TBEV and NTR controls. Four individual organoids per cell line per condition were combined into 9 pooled samples and analyzed. See also **Table B** for reference on a specific number of samples, replicates, and cell line details used in these experiments. **(B)** Volcano plots displaying DEGs (p.adj<0.1, log2FC>|0.5|) after HSV-1 (top) and TBEV (bottom) infection when compared to non-infected controls. The top 20 significantly altered genes are labelled with their gene name. **(C)** Top deregulated pathways enriched upon viral infection (HSV-1, TBEV) based on DEGs. **(D)** Targeted pathway analysis of DEGs (WT HSV-1-infected COs vs. corresponding non-infected controls) based on literature-defined AD-associated mechanisms ([40–43]; **Table E**).

To further assess whether HSV-1-induced transcriptomic changes align with pathways implicated in AD, we performed a targeted analysis. Based on the current literature [40–43], we selected ten key biological processes associated with AD onset and progression: (i) inflammation, (ii) oxidative stress and mitochondrial respiration, (iii) autophagy and mitophagy, (iv) loss of proteostasis, (v) cholesterol metabolism, (vi) cellular damage, (vii) senescence, (viii) intracellular communication, (ix) neurogenesis, and (x) amyloid processing (**Figure 4D, Table E**). Our analysis revealed that HSV-1 infection particularly affected genes involved in oxidative stress, ER stress, and apoptosis, findings that are consistent with the untargeted ORA results described above. Additionally, we identified deregulation in pathways related to endocytosis, amyloid metabolism, and cellular senescence. Together, these results underscore a fundamental difference in the molecular response of COs to HSV-1 and TBEV: while TBEV elicits a focused antiviral immune response, HSV-1 induces widespread transcriptional alterations across multiple cellular pathways with potential relevance to neurodegenerative disease.

### 3.5 Secretome Analysis Reveals Convergent Neurodegenerative and Senescence-associated Responses to HSV-1 and TBEV Infections

To complement our transcriptomic findings and to further explore the cellular response to viral infection, we analyzed the secretome of COs, focusing on proteins released into the extracellular space. Such secreted proteins could represent key mediators of intercellular communication and immune activation relevant to the pathogen hypothesis of neurodegeneration [6]. We performed an unbiased proteomic analysis of secreted proteins from D60 and D100 organoids derived from three independent iPSC lines, following the infection scheme presented in **Figure 2A**. Four individual organoids (=technical repetition) per cell line (WT1, 2, 3) per condition (NTR, HSV-1, TBEV), per timepoint (D60, D100; 74 individual samples) were measured and analyzed after technical tetraplicates were pooled (see **Supplementary Methods**, **Table B**). A list of differentially secreted proteins (DSPs) in HSV-1 and TBEV-infected COs as compared to their corresponding non-infected controls is included in **Table G**. Interestingly, in contrast to the transcriptomic data, principal component analysis revealed a greater similarity in secretory profiles between HSV-1 and TBEV-infected organoids (**Figure 5A**), despite HSV-1 infection leading to approximately twice as many deregulated proteins as TBEV (**Figure E.1**). Analysis of differential secretion as depicted in volcano plots (**Figure 5B**) revealed that most changes involved upregulated secreted proteins, and unbiased ORA of DSPs identified 12 commonly deregulated pathways across all datasets (**Figure 5C**). Notably, the most significantly affected pathways were again related to neurodegeneration, including those implicated in AD, Parkinson’s disease, amyotrophic lateral sclerosis, Huntington’s disease, and prion diseases (**Figure 5D**). This enrichment pattern was consistent in both younger (D60) and older (D100) organoids, regardless of the virus.

**Figure 5:**
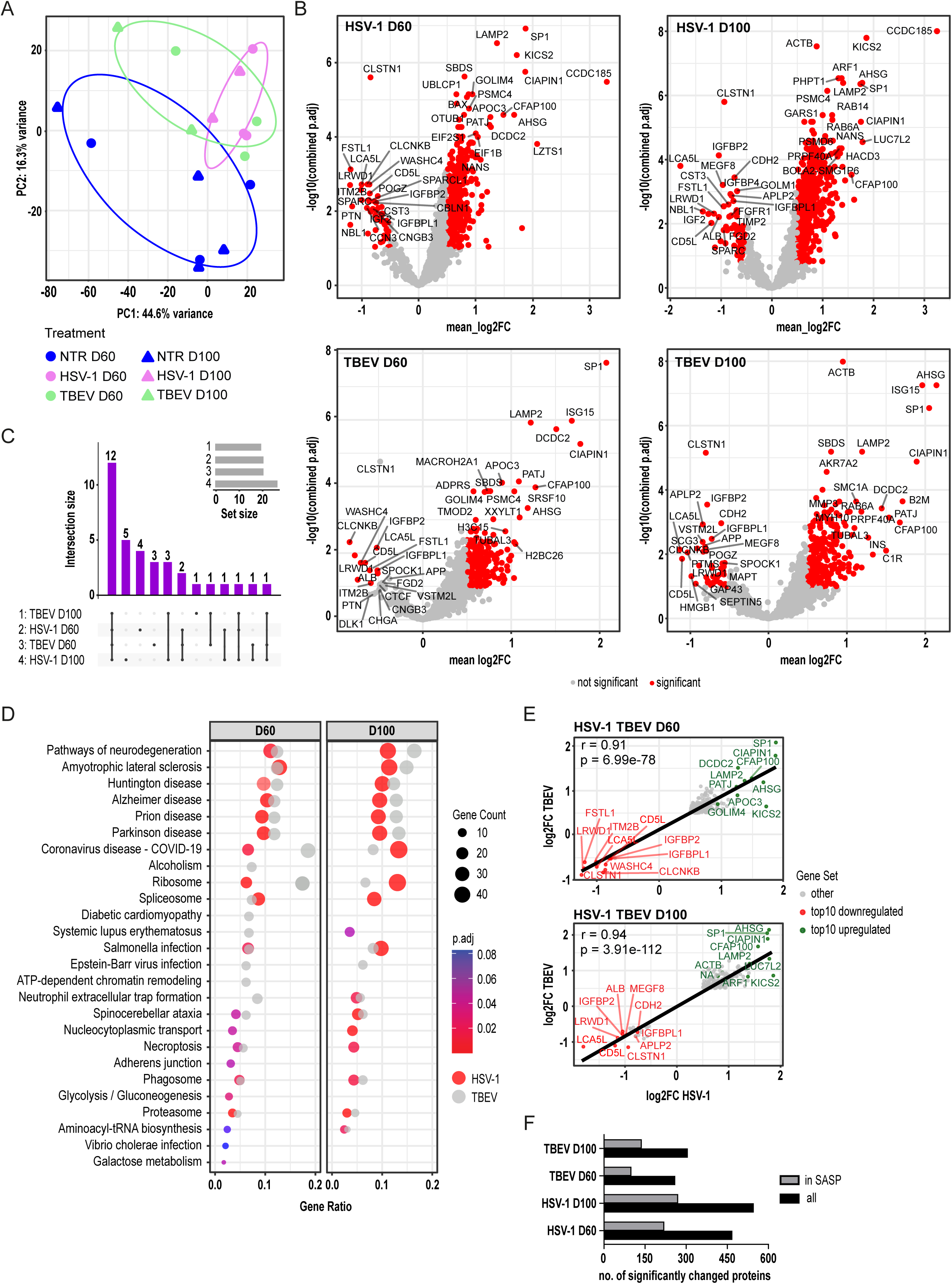
Secretome analysis reveals overlapping neurodegenerative and senescence-associated responses to HSV-1 and TBEV infections in COs. **(A)** Principal component analysis of proteins secreted into media from WT D60 and D100 COs following HSV-1 or TBEV infection and NTR controls. Four individual organoids (=technical repetition) per cell line per condition, per timepoint were measured separately and technical tetraplicates were pooled for analysis. See **Supplementary Methods** and also **Table B** for reference on a specific number of samples, replicates, and cell line details used in these experiments. **(B)** Volcano plots displaying DSPs (p.adj<0.1, log2FC>|0.5|) after HSV-1 (top) and TBEV (bottom) infection at D60 and D100 when compared to non-infected controls. The top 20 significantly altered genes are labelled with their gene name. **(C)** UpSet plot of pathways deregulated and shared/not shared across all datasets. **(D)** Top deregulated pathways (for each dataset) enriched upon viral infection based on DSPs (HSV-1 infection is shown in color, TBEV in greyscale). **(E–F)** DSPs (p.adj<0.1, log2FC>|0.5|) from the secretome datasets (HSV-1 and TBEV-infected COs vs. their non-infected controls) on D60 **(E)** and D100 **(F)** were plotted against each other, with the fitted line, Pearson’s correlation coefficient (r), and statistical significance. The top 10 DSPs (up/down, based on weighted scores) are labelled with protein/gene name. **(G)** Comparison of viral infection-induced secretome profiles with the SASP atlas [36].

Interestingly, when we compared DSPs that correlated between HSV-1 and TBEV infections at both D60 and D100 (Pearson’s correlation, r>0.9, p<0.05; **Figures 5E, F**), we observed a strong overlap in proteins associated with APP processing (e.g., SP1, CLSTN1, APLP2), autophagy (e.g., LAMP2, ARF1), IGF signaling (e.g., IGFBP2, IGFBPL1), and inflammation (e.g., CD5L, AHSG). These proteins displayed a consistent pattern of deregulation across both viral infections and time points. Additionally, we also noticed that the overlapping proteins at D60 were also associated with mitochondrial function and proteostasis, reflecting early neurodegenerative stress pathways. In contrast, the overlapping proteins at D100 shifted toward functions related to cell structure (e.g., ACTB), signaling, and systemic inflammation (e.g., ALB, AHSG), suggesting the emergence of broader stress and immune responses in more mature organoids.

Lastly, a comparison with the publicly available Senescence-Associated Secretory Phenotype (SASP) atlas [57] showed that approximately 50% of the secreted proteins following viral infection were SASP-related (**Figure 5G**). This proportion remained consistent across infection type and organoid age. Many of these SASP proteins were also part of neurodegenerative pathways identified in our ORA (**Figure E.2**), indicating that viral infection promotes a secretory profile enriched for senescence-associated and neurodegeneration-linked factors. Collectively, these findings demonstrate that even mild infections, such as those induced by TBEV, can activate secretory programs associated with neurodegeneration and cellular senescence, reinforcing the concept that pathogen exposure may contribute to long-term neuronal vulnerability and disease progression.

### 3.6 *PSEN1/2* Mutant Cerebral Organoids Exhibit Comparable Acute Molecular and Secretory Responses to Viral Infection as Wild-Type Counterparts

Thus far, for all our investigations, we have utilized three independent iPSC lines with WT variants of AD-related genes (*APP, PSEN1, PSEN2*, and *SORL1*). However, previous research using AD animal models has indicated that 5xFAD mice (expressing human *APP* and *PSEN1* with AD-linked mutations) exhibit a milder reaction to pathogen infections and survive longer, suggesting a potentially protective role of amyloids [49,58]. To investigate the effect of AD-linked mutations on the pathogen infection, we extended our analysis to iPSCs derived from patients with familial AD, carrying mutations in *PSEN1* (A246E) or *PSEN2* (I144N) (**Table A**). We previously demonstrated that this model accumulates large APP clusters over time [24]. Based on this, we hypothesized that such accumulation may alter cellular and molecular responses to infection.

Thus, following the same experimental approach as with WT lines, we differentiated three independent AD iPSC lines into COs and infected them with HSV-1 or TBEV and harvested them at both D60 and D100 **(Figure 6A)**. Immunofluorescent analyses of Aβ clusters (**Figure 6B**) and subsequent quantification of a total of 173 CO sections (**Figures 6C and F.1**) confirmed that *PSEN1/2* mutant organoids accumulate larger and more numerous Aβ clusters over time. However, similar to the WT organoids, we did not observe any infection-induced increase in Aβ clustering following HSV-1 or TBEV exposure. Consistently, a shift in the APP band on the Western blot was only detected after HSV-1 infection (**Figure 6D**), mirroring the response observed in WT organoids.

**Figure 6:**
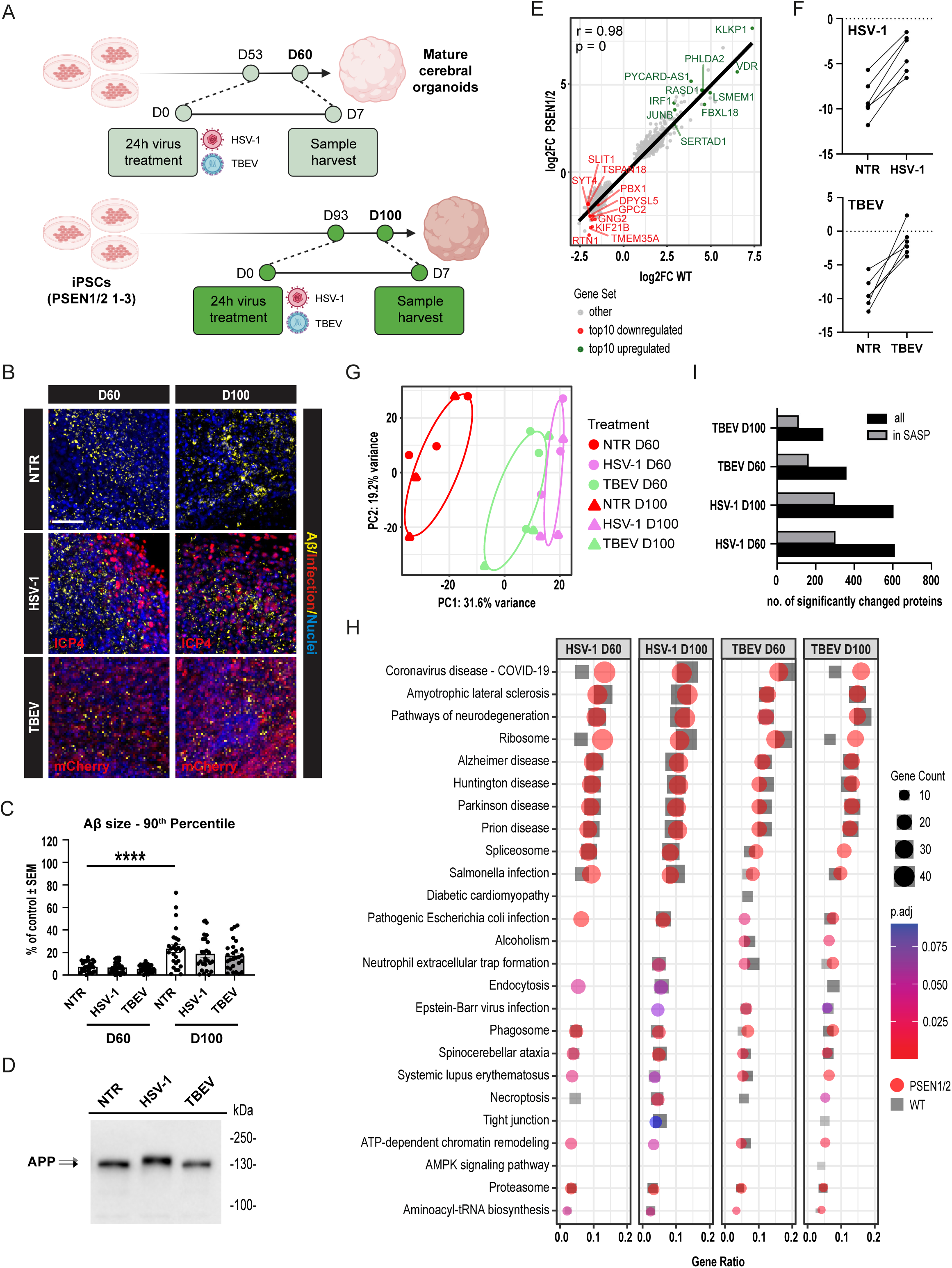
*PSEN1/2* mutant COs show similar acute molecular and secretory responses to viral infection as WT organoids. **(A)** Schematic overview of the experimental design for infection of *PSEN1/2* mutant COs with HSV-1 and TBEV at D60 and D100. **(B)** Representative immunofluorescence images showing Aβ accumulation in *PSEN1/2* mutant organoids at D60 and D100, with or without viral infection. Scale bar = 50 µm. **(C)** Quantification large Aβ clusters from images in **(B)** of a total of 173 CO sections. Data represent mean ± SEM. Statistical significance was determined by an unpaired t-test; ****p<0.0001; n≥3. **(D)** Representative Western blot analysis of full-length APP showing a band shift after HSV-1 infection. **(E)** DEGs (p.adj<0.1, log2FC>|0.5|) from the transcriptomic datasets comparing WT HSV-1 infected and *PSEN1/2* mutant HSV-1 infected COs (relative to their non-infected controls) were plotted against each other, with fitted line, Pearson’s correlation coefficient (r), and statistical significance. The top 10 DEGs (up/down, based on weighted scores) are labelled with gene names. **(F)** Senescence score calculated from transcriptomic data using a machine learning-based algorithm. **(G)** Principal component analysis of proteins secreted into media from *PSEN1/2* mutant D60 and D100 COs following HSV-1 or TBEV infection and NTR controls. Four individual organoids (=technical repetition) per cell line per condition per timepoint were measured (72 single COs) and technical tetraplicates were pooled for the analysis. (H) Top deregulated pathways enriched upon HSV-1 or TBEV infection based on DSPs (infection in PSEN1/2 mutant samples is shown in color, for WT in greyscale). (I) Proportion of secreted proteins matching entries in the SASP atlas after infection. See also **Table B** for reference on a specific number of samples, replicates, and cell line details used in these experiments.

We next analyzed the molecular response of *PSEN1/2* mutant organoids to viral infection using the same omics approaches as applied to WT organoids. Transcriptomic analysis of four individual organoids per cell line (*PSEN1/2* mutant 1, 2, 3) per condition (NTR, HSV-1, TBEV) combined into 9 pooled samples, indicated that HSV-1 infection had a more pronounced effect on D60 *PSEN1/2* mutant organoids compared to TBEV, as shown by Principal component analysis (**Figure F.2**). The significantly deregulated pathways overlapped with those previously identified in WT organoids (**Figure F.3**). Gene expression changes in AD organoids also correlated well with those in the WT, particularly in genes associated with the innate immune response to viral infection (Pearson’s correlation, r=0.98; **Figure 6E**). Consistent with our findings in WT organoids, targeted pathway analysis focusing on AD-related processes showed that *PSEN1/2* mutant organoids exhibited gene deregulation in pathways related to oxidative stress, ER stress, and senescence (**Figure F.4**). Interestingly, given the recurrent association with senescence, we further utilized transcriptomic data from both WT and *PSEN1/2* mutant organoids (a total of six datasets) to calculate a senescence score using a machine learning-based algorithm developed at the National Institute on Aging (Anerillas *et al*., in preparation). As shown in **Figure 6F**, these calculations confirmed that both HSV-1 and TBEV infections significantly increased the senescence score compared to pre-infection levels. These findings support the notion that cellular senescence may contribute to the host response to viral infection and could represent a mechanistic link in the Pathogen Infection Hypothesis of AD.

Importantly, secretome analysis of *PSEN1/2* mutant organoids revealed a similar pattern to that seen in WT lines. Principal component analysis of four individual organoids (=technical repetitions) per cell line (*PSEN1/2* mutant 1, 2, 3) per condition (NTR, HSV- 1, TBEV) per timepoint (D60, D100; 72 individual samples) confirmed that both HSV-1 and TBEV infections altered the secretory profile (**Figure 6G**). This profile again showed substantial overlap in pathway enrichment profiles, regardless of virus type. Unbiased ORA revealed shared enrichment in pathways associated with neurodegeneration, including AD, Parkinson’s, Huntington’s, and prion diseases (**Figure 6H**) across both time points. Notably, approximately 50% of the secreted proteins corresponded to components of the SASP atlas (**Figure 6I**), many of which were also implicated in the same neurodegenerative pathways identified in the overall secretome analysis (**Figure F.5**). In summary, our data demonstrate that COs derived from *PSEN1/2* mutant iPSC lines exhibit an acute molecular and secretory response to viral infection that closely resembles that of WT organoids. Whether the long-term consequences of infection differ between these genotypes remains an important question for future investigation.

### 3.7 Reanalysis of an Independent Dataset Confirms HSV-1-Induced Neurodegeneration and Senescence Pathways in Cerebral Organoids and Shows a Limited Protective Effect of Acyclovir

Finally, to cross-validate our findings, we reanalyzed publicly available transcriptomic data from Rybak-Wolf *et al.* [42], in which human iPSC-derived COs were used to model HSV-1-induced viral encephalitis (**Figure 7A**). This study reported significant disruption of tissue architecture, neuronal function, and cellular transcriptomes following HSV-1 infection and also employed 3’mRNA-seq for data analysis. Interestingly, functional experiments using acyclovir revealed that while this treatment effectively suppressed viral replication, it failed to prevent HSV-1-induced structural damage to neuronal processes and the neuroepithelium.

**Figure 7:**
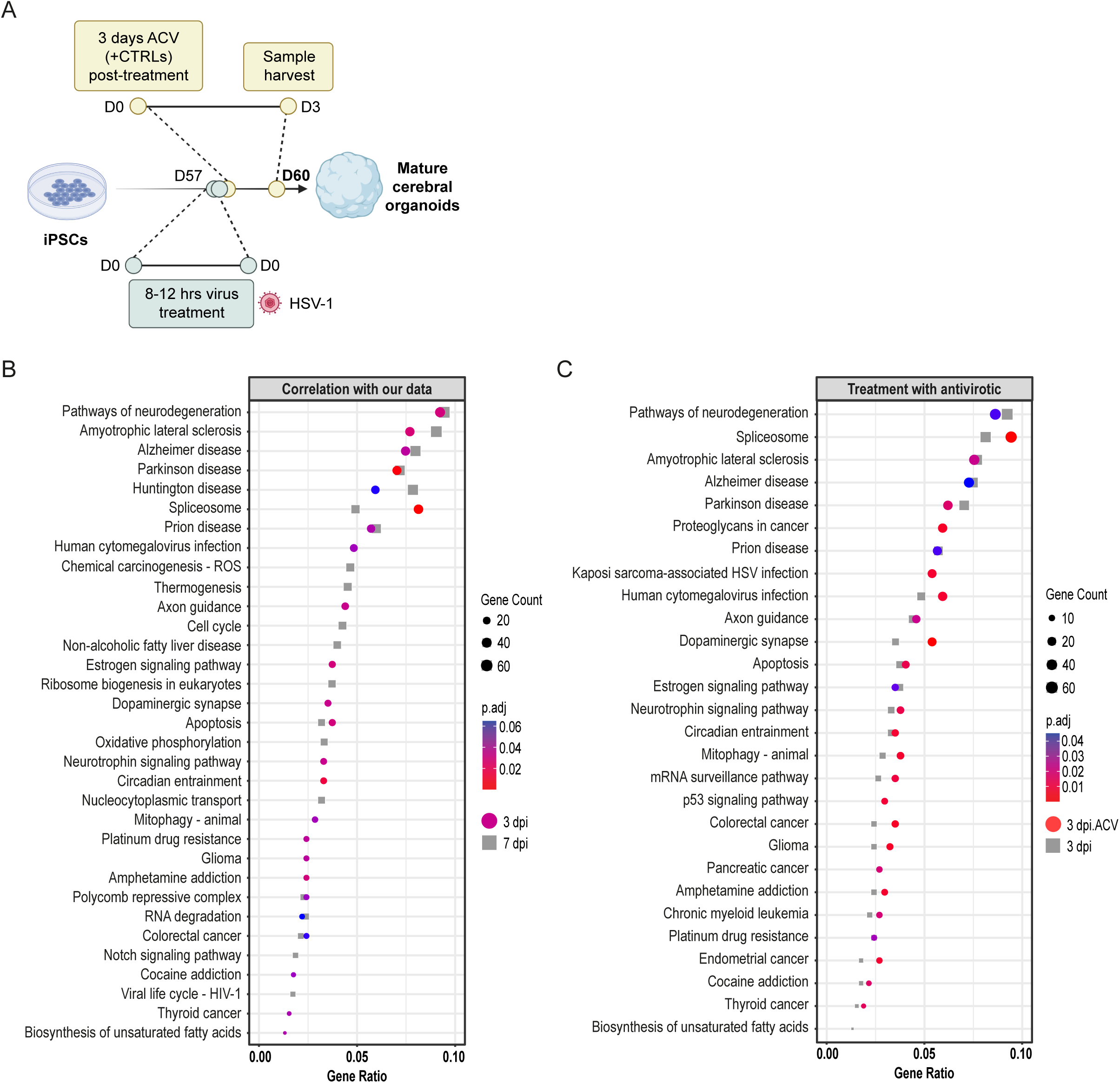
Reanalysis of independent transcriptomic data confirms HSV-1-induced activation of neurodegeneration and senescence pathways in COs, with limited protective effect of acyclovir. **(A)** Schematic representation of the experimental design from Rybak-Wolf *et al.* [42], in which human iPSC-derived COs were infected with HSV-1 and analyzed 3 dpi, with or without acyclovir treatment. 3’ mRNA-seq was used to assess transcriptomic changes. **(B)** Top deregulated pathways enriched upon HSV-1 infection, based on DEGs (3dpi vs. non-infected control) in D60 organoids from the Rybak-Wolf *et al.* [42] dataset. Pathways at 3 dpi are shown in color, while corresponding data from our 7 dpi dataset (Figure 4) are displayed in greyscale. **(C)** Top deregulated pathways enriched upon HSV-1 infection based on DEGs (D60 COs 3 dpi vs. D60 COs 3 dpi with acyclovir treatment; 3 dpi with acyclovir treatment shown in color, 3 dpi shown in greyscale).

Notably, the original study did not specifically investigate transcriptomic changes associated with neurodegeneration or cellular senescence. We, therefore, reanalyzed their 3’mRNA-seq data, focusing on D60 COs collected at 3dpi, both with and without acyclovir treatment. Using pathway enrichment analyses over GO and KEGG databases, we found that the datasets exhibited striking similarities to our own transcriptomic profiles. As depicted in **Figure 7B**, enriched KEGG terms included pathways related to neurodegeneration, such as AD, Parkinson’s disease, and prion diseases, consistent with our own findings from infected D60 organoids. When performing targeted pathway analysis using our pre-selected terms related to AD (**Table E**), we again identified strong enrichment in pathways linked to oxidative stress, ER stress, autophagy, apoptosis, synaptic transmission, exocytosis, and senescence (**Figure G.1**). These results closely mirror our targeted analyses in both WT and *PSEN1/2* mutant COs (see **Figures 4D** and **F.4**), further reinforcing the interpretation that HSV-1 infection activates transcriptional programs characteristic of early neurodegeneration.

Crucially, the availability of acyclovir-treated samples in this dataset allowed us to assess whether antiviral treatment modulated the expression of these pathways. As shown in **Figure 7C**, we observed no attenuation of neurodegeneration- or senescence-related pathway enrichment following acyclovir treatment, nor did it affect the targeted analysis and the pathways related to AD (**Figure G.2**). This result aligns with conclusions from the original study, which reported that acyclovir suppressed viral replication but did not prevent structural or transcriptomic damage. The authors further proposed that co-treatment with anti-inflammatory compounds, such as necrostatin-1 or bardoxolone methyl, was required to rescue infection-induced phenotypes. In summary, this reanalysis of an independent dataset robustly supports our findings, confirming that HSV-1 infection induces gene expression changes associated with neurodegeneration and cellular senescence, and further suggests that antiviral treatment alone may be insufficient to fully prevent these pathogenic responses in COs.

## 4. Discussion

The accumulation and aggregation of Aβ peptides have long been regarded as a hallmark of AD, yet the physiological function of these peptides and the triggers for their pathological deposition remain incompletely understood. The Pathogen Infection Hypothesis posits that Aβ may act as an evolutionarily conserved antimicrobial peptide, aggregating to entrap invading pathogens such as HSV-1 [6,47,49,59]. Importantly, this hypothesis also postulates that besides the entrapment of the evading pathogen(s), this infection also elicits a molecular response, which involves mainly the inflammatory response that may cause neuronal damage over time, but that also causes direct damage and cell death [6]. In this study, we systematically evaluated the response of human iPSC-derived 2D neuronal cultures and 3D COs to infection by HSV-1 or TBEV, with the goal of dissecting acute host cellular and molecular responses and their potential connection to AD-like pathology. Given the relevance of familial AD mutations for the accumulation of Aβ peptides, we extended our analyses to organoids derived from *PSEN1* (A246E) and *PSEN2* (I144N) mutant iPSC lines.

### 4.1 Protein Aggregation

One of the central components of the Pathogen Infection Hypothesis is protein aggregation as a form of innate immune defense [6,7]. In the context of AD, Aβ has been proposed to act as antimicrobial peptide, capable of opsonizing or neutralizing invading pathogens [6,47–49,60]. This hypothesis is supported by *in vivo* evidence from familial AD mouse models, in which Aβ-overproducing transgenic mice infected with HSV-1 exhibit increased survival compared to non-transgenic controls, suggesting a protective role for Aβ-mediated aggregation in response to infection [49]. However, the findings from other studies have not consistently supported these observations [17,18]. It was also unclear whether such mechanisms are active in human 3D brain-like systems. While several studies using neuronal cultures embedded in 3D scaffolds or matrices reported Aβ accumulation following HSV-1 infection, other reports using self-organizing iPSC- derived COs failed to observe similar APP clustering [51–53,60]. Moreover, it was unknown whether non-herpes neurotropic viruses, such as TBEV, can elicit comparable responses.

Our study now provides new evidence that HSV-1, but not TBEV, robustly induces Aβ clustering in 2D iPSC-derived neuronal cultures and 3D COs derived from both WT and *PSEN1/2* mutant iPSCs. Importantly, this clustering response required high concentrations of extracellular Aβ peptides, which could be achieved either through endogenous secretion in 2D neurons (and likely also in 3D models based on mature iPSC-derived neurons) or exogenous supplementation in 3D iPSC-derived self- organizing organoids, supporting the postulate that Aβ functions as a pathogen- responsive seeding molecule. These findings also strongly suggest that Aβ aggregation is not a default feature of neurotropic viral infections but is rather a specific and conditional response that depends on both pathogen type and peptide abundance. The lack of comparable responses to TBEV, despite robust infection and immune activation, further underscores the selectivity of Aβ antimicrobial activity and suggests that specific viral features (e.g., envelope structure or glycoprotein density) may be required for effective opsonization.

Interestingly, our study also uncovered a previously unreported post-translational modification of APP occurring exclusively after HSV-1 infection, observed as a consistent shift in the full-length APP band in Western blot analyses. This modification was independent of Aβ clustering or Aβ pre-treatment. The nature of this modification, whether it reflects altered phosphorylation, glycosylation, cleavage, or interaction with viral or host proteins, remains to be determined. Nonetheless, it indicates that HSV-1 may exert direct intracellular effects on APP processing beyond promoting extracellular aggregation.

### 4.2 Inflammation and Neuronal Damage

In addition to protein aggregation, the Pathogen Infection Hypothesis suggests that viral infections may trigger chronic inflammation or directly damage neural tissues, thereby initiating or exacerbating neurodegenerative processes [6]. Persistent neuroinflammation is a well-established hallmark of AD [61], and several pathogens have been shown to provoke sustained immune responses within the central nervous system [47,62]. In some cases, viruses can directly infect neurons or glial cells, leading to cellular dysfunction or death [63]. Notably, HSV-1 has been detected in the brains of AD patients and is known to infect neurons, raising the possibility of a direct contribution to disease progression. Moreover, recent population studies, such as the work by Eyting *et al.* [14], have suggested that zoster vaccination may reduce the risk or delay the onset of dementia, reinforcing a potential link between latent viral infections and neurodegenerative outcomes. Additionally, multiple studies have modeled HSV-1 encephalitis in human COs, demonstrating that the virus impairs neuroepithelial identity and damages neuronal architecture [20,42]. The Rybak-Wolf *et al.* [42] study further reported that acyclovir, while effective in halting HSV-1 replication, did not prevent tissue damage or transcriptomic disruption. Their transcriptomic analyses identified tumor necrosis factor signaling as a central driver of pathology, and only co-treatment with anti-inflammatory agents, such as necrostatin-1 or bardoxolone methyl, was able to restore neuronal integrity. Despite these findings, none of the existing studies directly investigated whether HSV-1 infection activates neurodegeneration-specific pathways, nor did they assess the response to other neurotropic viruses such as TBEV.

In our study, we now demonstrate that HSV-1 infection induced a broad and complex transcriptional response in both WT and *PSEN1/2* mutant COs, deregulating genes significantly involved in pathways related to neurodegeneration, including Alzheimer’s, Parkinson’s, and prion diseases. Targeted pathway analysis further identified hallmark AD-related processes, such as oxidative stress, ER stress, and apoptosis, as being transcriptionally upregulated in HSV-1-infected organoids. In contrast, TBEV infection induced a more restricted antiviral immune response, primarily through the activation of interferon-stimulated genes, without transcriptionally engaging broader neurodegenerative pathways. Importantly, despite such a mild reaction at a transcriptional level, TBEV elicited a relatively broad response at the secretome level, similar to that induced by HSV-1. This data suggests that even a mild viral infection could profoundly affect the secreted proteins, key mediators of intercellular communication and immune activation.

Importantly, we also validated our findings by reanalyzing publicly available RNA-seq data from Rybak-Wolf *et al.* [42]. Consistent with our results, their HSV-1-infected COs had robust activation of neurodegeneration-related pathways. Importantly, while acyclovir effectively halted viral replication, it failed to reverse the upregulation of these damaging transcriptional programs, mirroring the original observation of the study that neuronal damage persisted unless anti-inflammatory agents were also administered.

### 4.3 Cellular Senescence

One of our novel findings is the consistent induction of cellular senescence following viral infection in human COs. Cellular senescence is a state of permanent cell cycle arrest, typically accompanied by a distinct proinflammatory secretory profile known as the SASP [64]. While originally studied in the context of tumor suppression and aging, senescence has recently been implicated in the pathogenesis of neurodegenerative diseases, including AD [65,66]. In the AD brain, the accumulation of senescent glial and neuronal cells contributes to chronic inflammation, disrupted proteostasis, and tissue dysfunction, processes tightly linked to cognitive decline and disease progression. Interestingly, senolytics have already been shown to work in animal models of AD [67,68]. Importantly, viral infections may trigger senescence [69] either directly, through mechanisms such as DNA damage, or indirectly via prolonged exposure to cytokines like interferons and tumor necrosis factor-alpha. These signals can promote so-called “paracrine senescence” in nearby uninfected cells, contributing to broader tissue-level responses. Curiously, despite these theoretical links, the role of pathogen-induced senescence and its connection to neurodegenerative mechanisms have not been directly examined in human-relevant models.

Our study now addresses this gap and demonstrates that HSV-1-infected COs upregulate senescence-associated genes and pathways in both our dataset and a published dataset. Furthermore, we applied a machine learning-based methodology to calculate the “senescence score” (Anerillas *et al.*, in preparation) from six RNA-seq datasets (three WT and three *PSEN1/2* mutant organoids). Importantly, we found that HSV-1 infection consistently increased the senescence score compared to non-infected samples. Lastly, proteomic profiling of the secretome triggered by viral infections by both HSV-1 and TBEV further supported this, showing that approximately 50% of the deregulated proteins corresponded to entries in the SASP atlas. These findings suggest that senescence is not merely a byproduct of infection but a coordinated response, potentially playing a dual role in antiviral defense and the initiation of neurodegenerative cascades.

## 5. Summary

Our study provides experimental evidence that HSV-1, but not TBEV, induces Aβ clustering and activates transcriptional and secretory programs associated with neurodegeneration and cellular senescence in human iPSC-derived neuronal cultures and COs. These effects were consistent across WT and *PSEN1/2* mutant backgrounds and were validated by the reanalysis of independent datasets. Notably, these datasets also suggested that while antiviral treatment halted viral replication, it failed to reverse HSV-1-induced activation of neurodegenerative and senescence-related pathways. Taken together, these findings support a mechanistic link between viral infection and AD pathogenesis and underscore the need to consider infection-driven senescence and inflammation in the development of future therapeutic strategies.

## Acknowledgements

The authors would like to thank Prof. Ales Hampl for his continuous support. We are also grateful to Prof. Jiri Damborsky, Prof. Zbynek Prokop, and Prof. Robert Harris for their critical reading of the manuscript. We extend our heartfelt thanks to Dr. Pavel Abaffy and Dr. Zuzana Matusova for sharing their expert knowledge and valuable help. Finally, we thank Richard Zimmermann for his help with language editing.

We acknowledge the CF Genomics and the CF Bioinformatics supported by the NCMG research infrastructure (LM2023067 funded by MEYS CR) for their support with obtaining the genomic data presented in this paper and the support of the GeneCore Facility at the Institute of Biotechnology of the Czech Academy of Sciences for help with the initial data processing. We also thank the CELLIM core facility, supported by the Czech-BioImaging large research infrastructure project (LM2023050 funded by MEYS CR), for their support in acquiring imaging data used in this work.

## 10. Consent Statement

Not applicable. The human-derived iPSCs used in this study were obtained from previously published sources, where informed consent was appropriately addressed.

## 9. Appendices

### Supplementary Methods

#### Supplementary Figures

Figure A.1, A.2

Figure B

Figure C

Figure D

Figure E.1, E.2

Figure F.1, F.2, F.3, F.4, F.5

Figure G.1, G.2

#### Supplementary Tables

Table A: Cell lines used

Table B: Number of samples, replicates, and cell line details from all experiments performed in this study

Table C: Antibodies Table D: PCR Primers

Table E: List of selected pathways connected to neurodegeneration

Table F: List of deregulated/measured genes expressed in HSV-1- and TBEV-infected cerebral organoids.

Table G: List of deregulated/measured secreted proteins in HSV-1- and TBEV-infected cerebral organoids.

