## Supplementary_Figure_A for "Viral Infection Induces Alzheimer’s Disease-Related Pathways and Senescence in iPSC-Derived Neuronal Models"

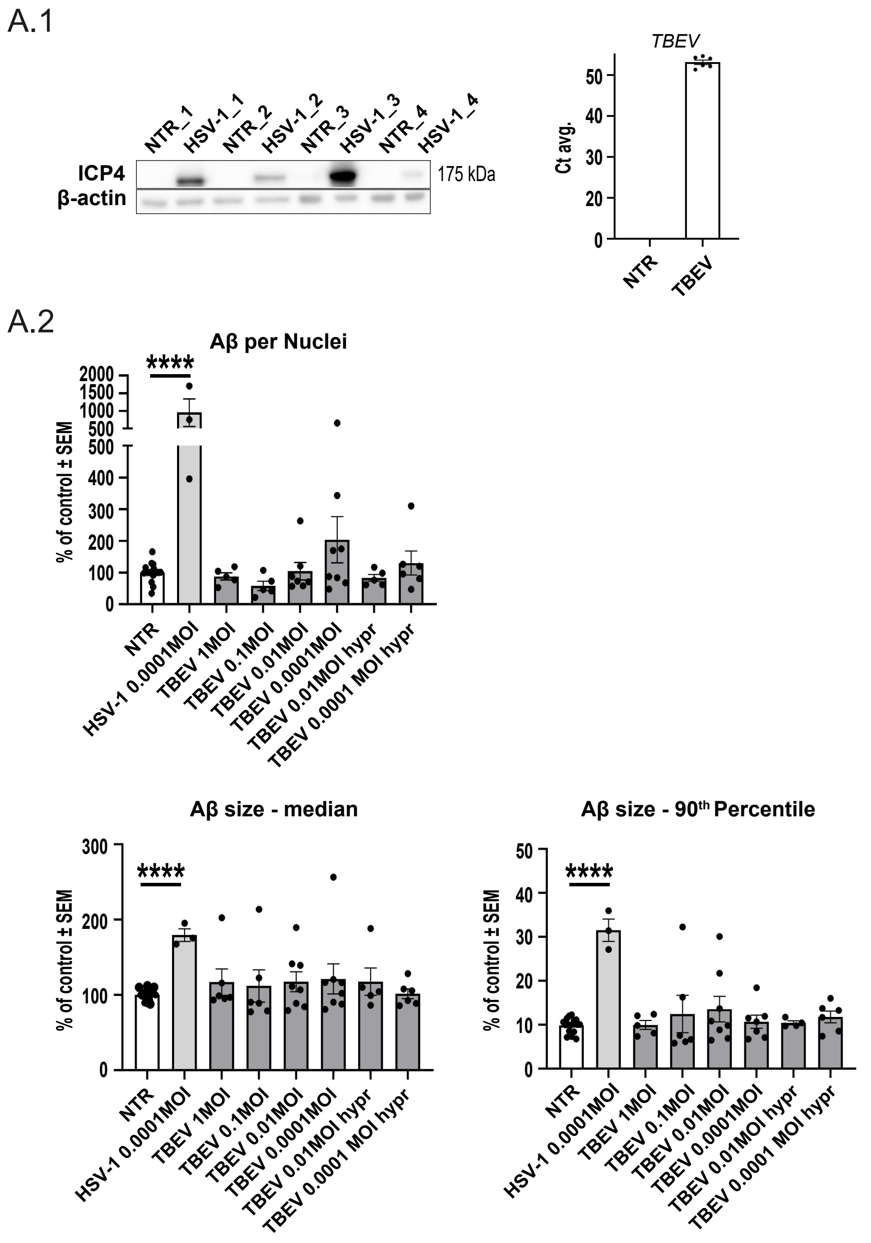


**Figure A: Differential accumulation of Aβ in 2D neuronal cultures following HSV-1 and TBEV infections.**

**Figure A: Differential accumulation of Aβ in 2D neuronal cultures following HSV-1 and TBEV infections. (A.1)** Validation of viral infection by Western blotting and qRT-PCR. For HSV-1, four independent infections were evaluated for the presence of ICP4. For TBEV detection, we used qRT-PCR, which was designed to detect a specific site of TBEV genomic RNA. NTR neurons were used as a control and Ct average (CT avg.) is plotted due to non-detectable expression in NTRs. **(A.2)** Image analysis and quantification of Aβ signal after infection by TBEV with different MOIs or TBEV strains. Aβ signal with every MOI for TBEV was compared to NTR control neurons and to neurons infected with HSV-1 at MOI 0.0001. We assessed i) volume of Aβ particles normalized to the volume of cell nuclei per image; ii) median volume of Aβ; and iii) $90^{th}$ percentile of Aβ particle size. Each dot represents one biological replicate, n≥3, significance was evaluated using unpaired t-test, error bars represent mean ±SEM; ****p<0.001. See **Table B** for reference on specific number of samples, replicates and cell line details.
