## Supplementary_Figure_B for "Viral Infection Induces Alzheimer’s Disease-Related Pathways and Senescence in iPSC-Derived Neuronal Models"

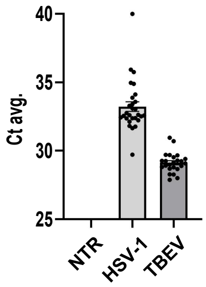


**Figure B: Confirmation of viral infection in COs.**

**Figure B: Confirmation of viral infection in COs.** qRT-PCR validation of HSV-1 and TBEV infection in COs harvested at D60. Ct average (CT avg.) is plotted due to non-detectable expression in NTRs. Each dot represents one biological replicate, $n \geq 3$. See **Table B** for reference on specific number of samples, replicates and cell line details.
