## Supplementary_Figure_C for "Viral Infection Induces Alzheimer’s Disease-Related Pathways and Senescence in iPSC-Derived Neuronal Models"

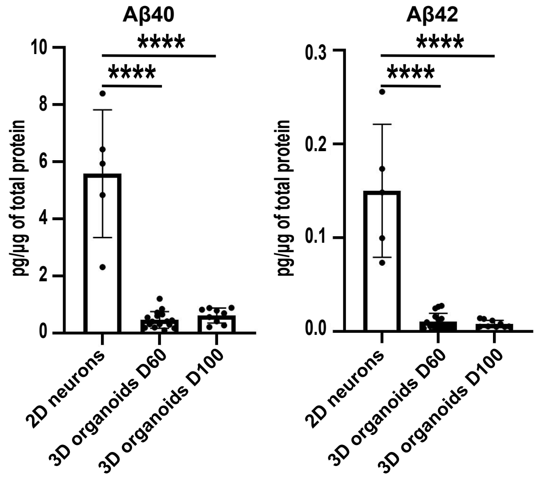


**Figure C: Comparison of secreted Aβ peptide levels in 2D neuronal cultures and 3D COs.**

**Figure C: Comparison of secreted Aβ peptide levels in 2D neuronal cultures and 3D COs**. Quantifying Aβ40 and Aβ42 levels in the culture medium of 2D neuronal cultures and 3D COs measured by ELISA. Data represent mean ± SEM from $n \geq 4$ of independent experiments. Statistical analysis was performed using unpaired t-test; ****$p < 0.001$. See **Table B** for reference on specific number of samples, replicates and cell line details.
