## Supplementary_Figure_D for "Viral Infection Induces Alzheimer’s Disease-Related Pathways and Senescence in iPSC-Derived Neuronal Models"

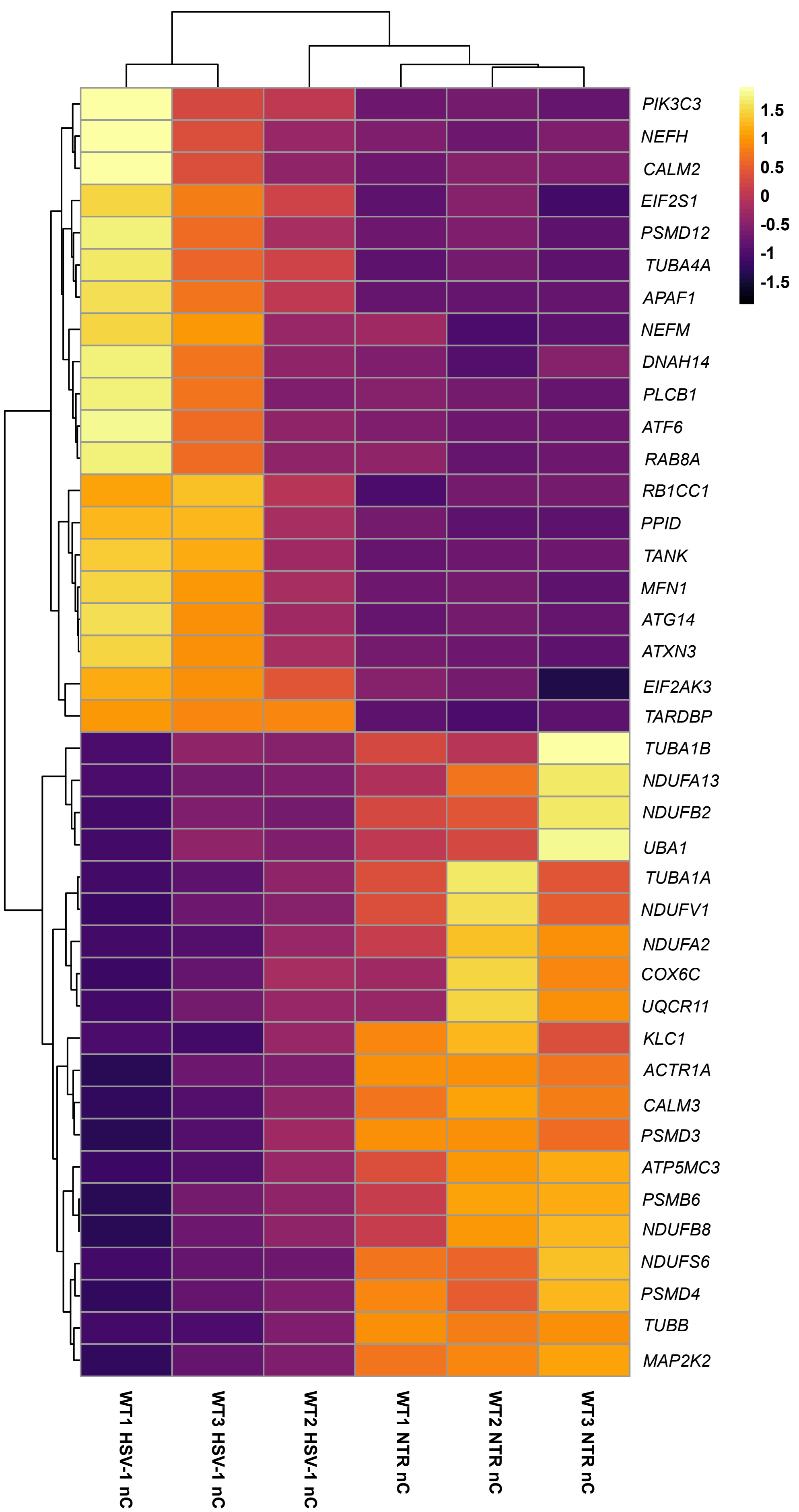


**Figure D: HSV-1-induced transcriptional changes affect genes involved in neurodegeneration-associated pathways.**

**Figure D: HSV-1-induced transcriptional changes affect genes involved in neurodegeneration-associated pathways.** Heatmap displaying the top 20 most upregulated and top 20 most downregulated genes from HSV-1-infected COs when compared to non-infected controls (p.adj<0.5, log2FC>|0.5|, top 40 DEGs selected based on their weighted scores).
