## Supplementary_Figure_E for "Viral Infection Induces Alzheimer’s Disease-Related Pathways and Senescence in iPSC-Derived Neuronal Models"

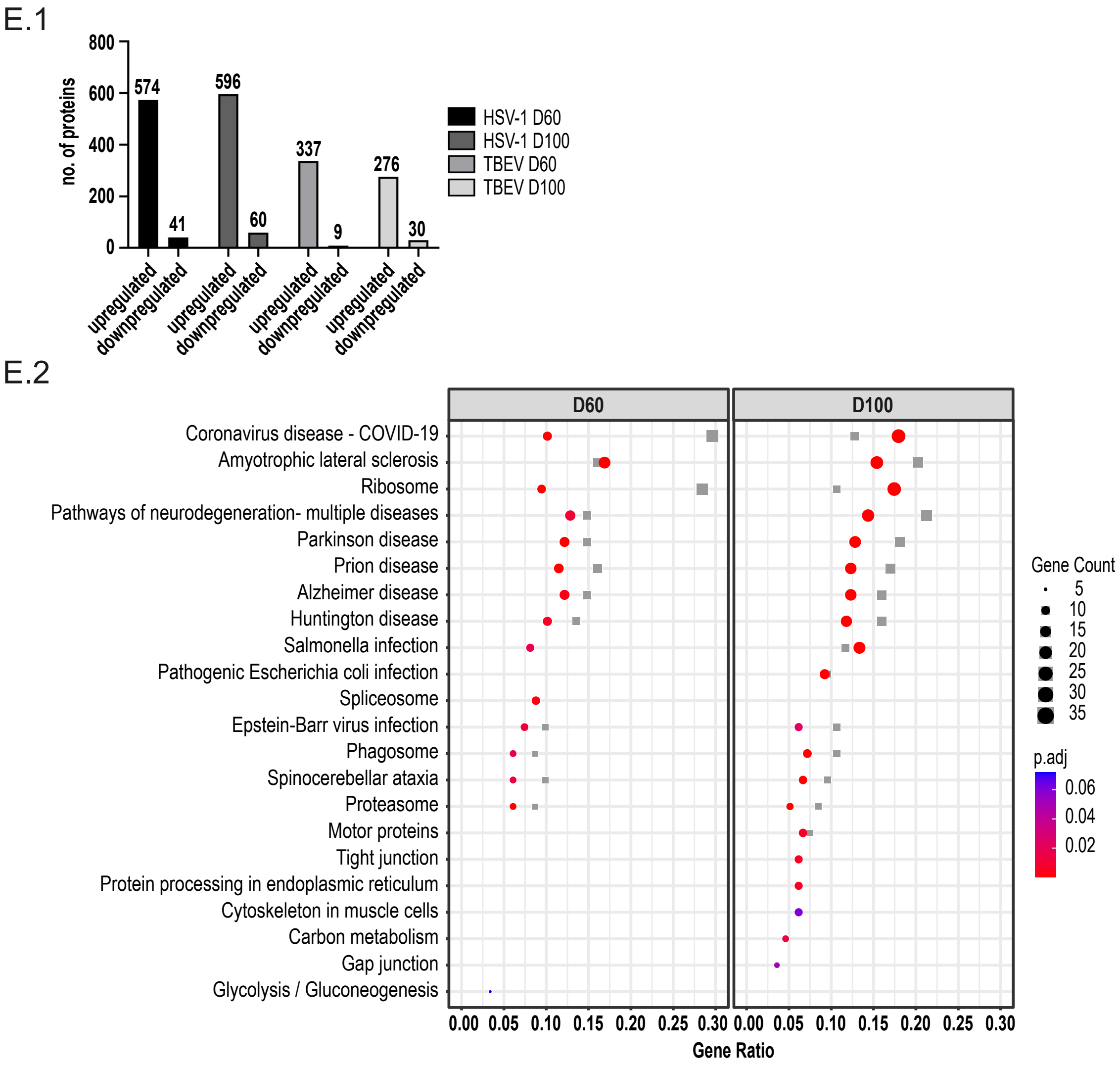


**Figure E: Analyses of secretome deregulation in viral-infected COs.**

**Figure E: Analyses of secretome deregulation in viral-infected COs. (E.1)** Bar plot comparing the number of significantly deregulated secreted proteins in D60 and D100 organoids following HSV-1 and TBEV infection. **(E.2)** Top deregulated pathways enriched upon viral infection (HSV-1 and TBEV) based on DSPs related to SASP (HSV-1 infection is shown in color, TBEV in grey scale).
