## Supplementary_Figure_F for "Viral Infection Induces Alzheimer’s Disease-Related Pathways and Senescence in iPSC-Derived Neuronal Models"

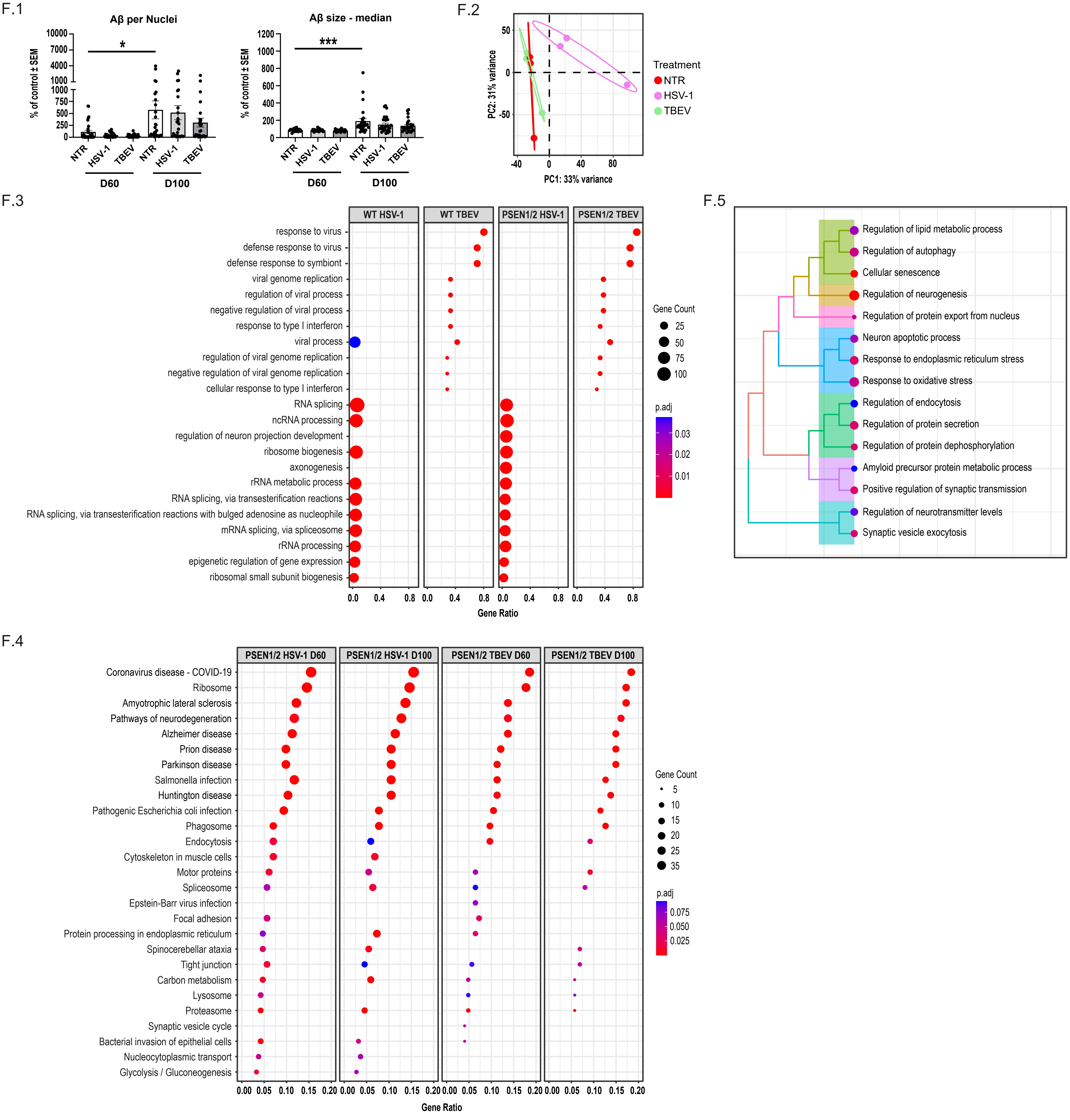


**Figure F: Transcriptomic and secretome profiles of PSEN1/2 mutant COs after HSV-1 and TBEV infection.**

**Figure F: Transcriptomic and secretome profiles of *PSEN1/2* mutant COs after HSV-1 and TBEV infection. (F.1)** Quantification of Aβ clusters in *PSEN1/2* mutant organoids at D60 and D100. Data represents analysis of mean ± SEM of a total of 173 CO sections. Statistical significance was determined by an unpaired t-test; ***$p<0.001$, *$p<0.1$; $n\geq3$. **(F.2)** Principal component analysis of transcriptomic data from *PSEN1/2* mutant iPSC-derived COs harvested at D60 following 7 dpi with HSV-1 or TBEV and NTR controls. Analysis was performed on 4 individual organoids per cell line per condition, combined into 9 pooled samples. **(F.3)** Top deregulated pathways enriched upon viral infection (HSV-1, TBEV) based on DEGs. **(F.4)** Targeted pathway analysis of DEGs (*PSEN1/2* mutant HSV-1 infected COs vs. corresponding non-infected controls) based on literature-defined AD-associated mechanisms ([40–43]; **Table E**) **(F.5)** Top deregulated pathways enriched upon viral infection (HSV-1, TBEV) based on DSPs secreted from *PSEN1/2* mutant COs related to SASP. See also **Table B** for reference on a specific number of samples, replicates, and cell line details used in these experiments.
