## Supplementary_Figure_G for "Viral Infection Induces Alzheimer’s Disease-Related Pathways and Senescence in iPSC-Derived Neuronal Models"

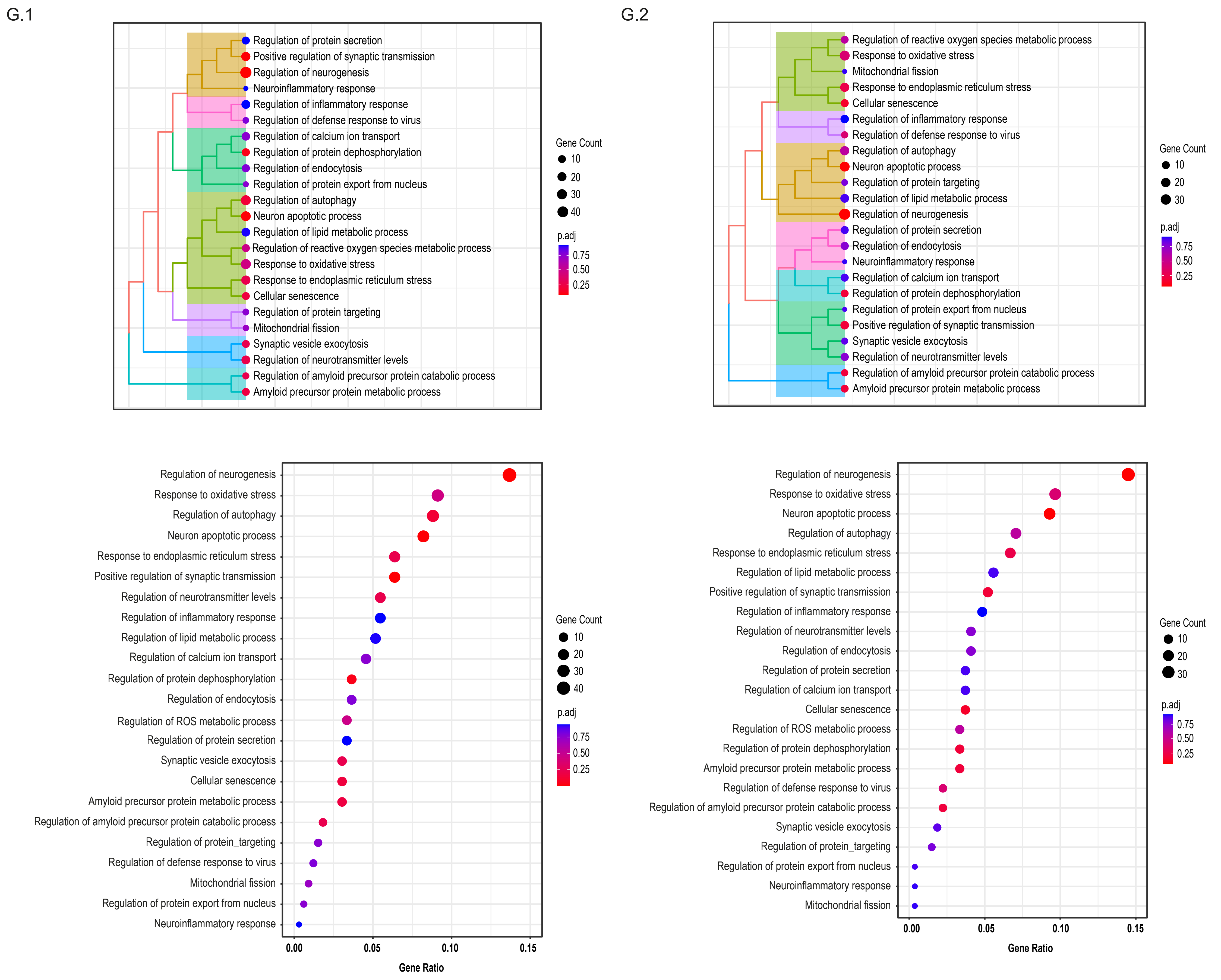


**Figure G: Targeted pathway analysis of HSV-1-infected COs from Rybak-Wolf et al.**

**Figure G: Targeted pathway analysis of HSV-1-infected COs from Rybak-Wolf *et al.*** [42] **(G.1)** Targeted analysis of DEGs in between HSV-1-infected COs and their corresponding non-infected controls with emphasis on literature-defined AD-associated mechanisms ([40–43]; **Table E**) showing strong enrichment in oxidative stress, ER stress, autophagy, apoptosis, synaptic transmission, exocytosis, and senescence—closely matching our analyses in WT and *PSEN1/2* mutant organoids (**Figures 4D and F.4**). **(G.2)** Comparison of targeted AD-related pathway enrichment in acyclovir-treated and untreated conditions. The top charts show clustered data, and the bottom charts show the same data sorted based on p.adj and gene ratio.
