## Supplementary_Methods for "Viral Infection Induces Alzheimer’s Disease-Related Pathways and Senescence in iPSC-Derived Neuronal Models"

1. **Immunochemistry and microscopic analysis**
   1.1 **Immunocytochemistry of 2D neurons and microscopy**

Immunocytochemistry was performed as described previously [1]. Briefly, cells were fixed with 4% paraformaldehyde, permeabilized using 0.2% Triton X100 in 1xPBS for 15 min, and incubated with primary antibodies overnight at 4 °C. Secondary antibodies and DAPI (D9542, Merck) were diluted in the permeabilization buffer and incubated with cells for 1 h at room temperature. After incubation, the slides were washed carefully with PBS, dried, and mounted onto microscopic slides with Mowiol Reagent (Merck). Used antibodies can be found in **Table C**.

Samples were imaged with the widefield microscope Zeiss Axio Imager.Z1, equipped with a ZEISS Colibri 7 and EC Plan-Neofluar 40x / 0.75 AIR objective using ZEN Blue software (Zeiss). Hoechst was detected using UV LED with illumination wavelength 385/30 nm, excitation filter 375/395 nm and emission filter 410-440 nm. AF568 was detected using green LED with illumination wavelength 555/30 nm, excitation filter 559/585 nm and emission filter 600-690 nm. Images with 0.325 × 0.325 × 1 μm pixel size were acquired using a monochromatic camera Hamamatsu ORCA Fusion (sCMOS sensor). A total of 49 tiles were acquired from each sample and subsequently analyzed for Aβ accumulation.

1.2 **Preparation of 3D cerebral organoid thick sections, immunohistochemistry, glycerol clearing and microscopy**

Before sectioning of cerebral organoids (COs) on the vibratome and immunostaining methods, harvested COs were fixed with 3.7% paraformaldehyde for one hour [1].

**Vibratome:** Fixed organoids were embedded in 4% agarose gel solution (A9414, Merck) and cut into 250-μm-thick sections with the Leica VT1000 S vibratome (Leica Biosystems) using a speed of 0.075 mm/s and 100 Hz frequency. For subsequent immunohistochemistry staining, three COs per cell line/condition/timepoint were selected. Additionally, three thick sections from each CO were chosen to be analyzed.

**Immunohistochemistry and glycerol clearing:** CO thick sections were permeabilized and blocked in blocking buffer (0.5% Triton X100, 5% BSA in PBS) for 5 hours. Sections were incubated with primary antibodies diluted in the blocking solution at 4 °C for 5 days and then with secondary antibodies at 4 °C overnight. Nuclei were visualized by Hoechst (H1399, Thermo Fisher Scientific). Labeled CO sections were cleared using 60% glycerol and 2.5M fructose at room temperature overnight. Used antibodies can be found in	**Table C**.

**Microscopy:** Samples of CO thick sections were imaged with the inverted microscope Zeiss Axio Observer.Z1 with confocal unit LSM 800, equipped with solid state lasers (405, 488, 561, 640 nm), using Plan-Neofluar 5x/0.16 AIR objective, Plan-Neofluar 10x/0.30 AIR and ZEN Blue software (Zeiss). Images with 0.777 x 0.777 x 20 µm (5x) and 0.624 x 0.624 x 6 µm (10x) pixel size were acquired using GaAsp PMT detectors. The acquisition parameters of detectors for Alexa Fluor 488, 568, and 647 were: 497‑553 nm, 574‑617 nm, and 656‑700 nm (emission wavelength range) and 1 µs (pixel dwell time). Scan mode was set up to frame, and the pinhole was set to 1AU. Line average of 2 was applied to all channels.

1.3 **Image analysis**

To evaluate Aβ accumulation in human neurons and CO thick sections, we used commercially available software Imaris version 9.8.2 (Bitplane). The detection of individual Aβ plaques was performed in Imaris using the “Surface” module. The estimated parameters included the volume of Aβ and nuclei. Parameters were automatically quantified using the Imaris software. Data was analyzed and plotted using GraphPad Prism version 9.

2. **qRT-PCR**

Viruses in the harvested COs were inactivated with 30 minutes exposure to the UV light, COs were then washed with PBS, lysed with 1 ml RNA Blue reagent (Top-Bio), and stored at -80°C. Total RNA was isolated separately from single COs in three or four biological replicates per all tested variants (non-treated=NTR, HSV-1, TBEV) with a Direct-zol RNA Microprep kit (ZymoResearch) according to the manufacturer’s instructions. NanoDrop 1000 (Thermo Fisher Scientific) was used for determining RNA concentration and purity. The isolated RNA was transcribed to cDNA using 200 ng of total RNA and Transcriptor First Strand cDNA Synthesis Kit (Roche) according to the manufacturer’s instructions with either random hexamers or oligo(dT). qRT-PCR was performed from the cDNA samples using LightCycler® 480 SYBR Green I Master kit (Roche) in 10 μl reaction on LightCycler 480 II (Roche) following optimized protocol: preincubation (95 °C for 5 min), followed by 45 cycles of amplification and detection (95 °C for 10 s, 60 °C for 10 s and 72 °C for 10 s). Ct values were calculated using the automated Second Derivative Maximum Method in LC480 software (Roche). Average Ct values are presented in the figures as the specific viral genes/gDNA were not detected in control samples. Specific primers are listed in **Table D** based on the literature [2,3].

3. **Western blotting**

Protein lysis and Western blot were performed as described previously (Fedorova et al., 2019). Briefly, protein samples were lysed in 1% SDS-lysis buffer, concentration was measured with DC Protein Assay Reagents Package (5000116, Bio-Rad) mixed with 10x Laemmli buffer and incubated at 95°C for 10 min. Proteins were separated on 10% or 15% Acrylamide gel and transferred onto PVDF membranes (Merck). Membranes were blocked and incubated with antibodies in 5% skimmed milk or BSA. Antibodies and specific conditions are listed in **Table C**. The results were visualized via ECL™ Prime (Amersham) using ChemiDoc™ Touch Imaging System (Bio-Rad).

4. **ELISA**

For ELISA, COs were moved into poly-HEMA-coated 12-well plates (1 CO per well) with 1.5 ml of Essential 6™ Medium (A1516401, Thermo Fisher Scientific). After 72 hours, the medium and the corresponding COs were collected and stored at −80 °C. The amount of Aβ40 and Aβ42 peptides in cell culture media was measured with Amyloid beta 40 Human ELISA Kit (KHB3481, Thermo Fisher Scientific) and Amyloid beta 42 Human ELISA Kit, Ultrasensitive (KHB3544, Thermo Fisher Scientific) according to the manufacturer’s instructions. For each condition, media were collected from four COs and all the samples were analyzed in technical duplicates.

To compare the amounts of Aβ40 and Aβ42 peptides from COs of different sizes, we measured the total protein concentration of lysed COs and used it for normalization. Organoids were lysed at 4 °C for 30 min in 50-100 μl of lysis buffer (50 mM Tris-HCl pH 6,8, 1% sodium dodecyl sulfate, 10% glycerol). Brief sonication was used to facilitate the lysis. The lysate was pelleted by centrifugation, and the clear supernatant was taken for protein concentration measurement using DC Protein Assay Reagents Package.

5. **RNA isolation and 3’mRNA-sequencing**

Viruses in the harvested COs were inactivated with 30 minutes exposure to the UV light, COs were then washed with PBS, lysed with 1 ml RNA Blue reagent (Top-Bio), and stored at -80 °C. Total RNA was isolated separately from single COs in 3-4 biological replicates from the same condition (e.g., WT1_1, WT1_2, WT1_3, WT1_4) that were finally equimolarly pooled to provide a pooled sample (e.g. WT1.NTR), this was done on six different sets of cell lines (WT1, WT2, WT3; PSEN1/2 mutant 1, 2, 3) in all experimental conditions (NTR, HSV-1, TBEV) with Direct-zol RNA Microprep kit (ZymoResearch) according to manufacturer’s instructions. RNA quality was assessed by TapeStation 2200 (Agilent Technologies; RNA Screen Tape), and only samples with RINe values ≥ 7.5 were used for library preparation. Poly-A selected libraries were made from 250 ng of total RNA using QuantSeq FWD 3’mRNA Library Prep Kit (Lexogen) in combination with UMI Second Strand Synthesis Module for QuantSeq FWD and Lexogen i5 6nt Unique Dual Indexing Add-on Kit (Lexogen) according to the manufacturer’s instructions. Quality control of library quantity and size distribution was done using QuantiFluor dsDNA System (Promega) and High Sensitivity NGS Fragment Analysis Kit (Agilent Technologies). The final library pool was sequenced on NextSeq 500 (Illumina) using High Output Kit v2.5 75 Cycles in single-end mode, resulting in an average of 10 million reads per sample.

5.1 **3’mRNA-seq Data Analyses and overrepresentation analyses**

Data analysis was performed in R statistical environment v4.4.2 [4]. The quality of raw reads was verified using FastQC v0.11.9, and the potential contamination was screened by FastQ_Screen v0.11.1 [5]. The low-quality reads, and adaptor sequences were removed using TrimmomaticSE v0.36 with parameters “HEADCROP:10 ILLUMINACLIP: Lexogen_quantseq.fa:2:30:10 LEADING:3 TRAILING:3 SLIDINGWINDOW:4:15 MINLEN:36”. Ribosomal and mitochondrial reads were removed using sortmerna v2.1b. BAM files with alignment were created with STAR v2.7.0 [reference Homo sapiens genome version GRCh38; [6] The count tables were generated using the script htseq-count v0.11.4 [7] with annotation version GRCh38.87 and parameter “-m union”. ENSEMBL-IDs were used as identifiers of transcripts. The counted data were subsequently analyzed using R-package DESeq2 v1.46.0 [8]. Rlog transformed data were processed in the principal component analysis. Differentially expressed genes (DEGs) were identified by the command ‘DESEQ’ with default parameters. ENSEMBL-IDs were converted into Gene symbol using org.Hs.eg.db v3.20.0 database [9]. DEGs (p.adj<0.1, |log2FC|>0.5, baseMean>100) were plotted with ggplot2 v3.5.1 [10] as volcano plots, gene names description was assigned to the top 20 DEGs (up/down) based on their weighted score (log2FC*(-log10(p.adjust)). Functional Gene-ontology overrepresentation analysis (ORA) was conducted using clusterProfiler package v4.14.3 [11,12] by enrichGO (*OrgDb = org.Hs.eg.db, ont = ‘BP’, pAdjustMethod = ‘BH’, pvalueCut off = 0.1, minGSSize = 10”)* or enrichKEGG command; pathways where at least 5 genes were overlapping were plotted using ggplot2 package v3.5.1 [10] or UpSetR v1.4.0 [13]. Heatmap of 40 top differentially expressed genes (based on weighted score log2FC*(-log10(p.adj)) from a KEGG pathway hsa05022 (Pathways of neurodegeneration - multiple diseases) [14] as plotted from normalized counts with pheatmap v.1.0.12 [15]. The correlation between DEGs and/or DSPs (decsribed in the MS section of the Methods) was counted with Pearson’s correlation of log2FC and plotted with ggplot2 v3.5.1 [6]. Targeted analyses were based on preselected GOBP (msigdbr package v7.5.1 [16] terms connected to Alzheimer's and HSV-1 infection [17–20] (**Table E**) using enricher() function from the clusterProfiler package v4.14.3 [7, 8] ORA was performed using the list of DEGs, with custom TERM2GENE and TERM2NAME mappings. The significance thresholds for p-value and q-value were set to 0.99 to ensure comprehensive term inclusion in this exploratory analysis. These terms were further clustered using pairwise_termsim() from the GOSemSim v2.32.0 package [21,22] and plotted with the treeplot() from enrichplot package v1.26.2 [23].

6. **Mass Spectrometry analysis of secretome and intracellular proteins**
   6.1 **Extraction of proteins from collected organoid E6 cultivation medium**

Cultivation E6 medium was collected from each CO (1 300 – 1 500 μl) and secreted proteins were extracted by protein precipitation after the addition of 1/100 of the total sample volume of 2% SDC and 1/10 of total sample volume of trichloroacetic acid (TCA). Protein precipitation was ongoing during 30 min (addition of SDC) and, subsequently, 90 min on ice (addition of TCA). The supernatants were removed from pellets after sample centrifugation at 14 000 g, 15 °C, for 30 min. Five hundred µl of pre-chilled acetone (-20°C) were added and protein precipitation was allowed to proceed at -20°C overnight. Supernatants were removed from pellets after sample centrifugation at 14 000 g, 15 °C, for 10 min. The possible remnants of acetone were removed by evaporation for 2 min and the pellets were re-dissolved in 50 μl of 50 mM TEAB. Total protein concentration was determined in all samples using microBCA protein assay (Thermo Fisher Scientific).

6.2 **Protein digestion**

Fifteen μg of proteins from each sample was taken. Also, 2 μg from each sample was combined together to design global internal standard (GIS). Enzymatic digestion of proteins was processed in 100 mM (TEAB). Disulfide bonds were reduced with 5 mM tris(2-carboxyethyl) phosphine hydrochloride at 37°C for 20 min, and free thiol groups were blocked using 10 mM methyl methanethiosulfonate at room temperature for 15 min. To remove of contaminant reagents, 6 volumes of (~360 µl) of pre-chilled acetone were added, and protein precipitation was allowed to proceed at -20 °C overnight. Samples were centrifuged at 12 000 g at 4 °C for 10 min and acetone was decanted without disturbing the pellet. The pellet was allowed to dry for 2 min to remove the acetone remnants and re-dissolved in 32.5 μl of 100 mM TEAB. Finally, proteins were digested by a mixture of lysyl endopeptidase and trypsin (Promega) at 1:25 enzyme to protein ratio (w/w) at 37 °C overnight.

6.3 **Tandem Mass Tag isobaric labeling of peptides**

After digestion, total peptide concentration was determined using Fluorometric Peptide Assay (Thermo Fisher Scientific) according to the manufacturer’s instruction. Tandem Mass Tag (TMT) 16plex (Thermo Fisher Scientific) isobaric label reagents were dissolved in 31 μl of acetonitrile (AcN) for 5 min with occasional vortexing. TMT tubes were briefly centrifuged to gather the solution. Volume of samples was adjusted to 33 µl using 100 mM TEAB and 15 µl of TMT channels (250 μg) were added to appropriate samples to design 10 multiplexes of 16 TMT channels. The volume of GIS was split into five equal aliquots (50 μl). Five aliquots of the last TMT channel 134 were re-dissolved in 20 μl of AcN (500 μg) and added to GIS aliquots. Isobaric labeling of peptides was allowed to proceed at room temperature for 60 min and the reaction was terminated by 5% hydroxylamine according to the manufacturer’s instructions. Fifteen different samples (~3.5 µg, 12 µg) and one GIS aliquot (5.7 µg, 15 µg) were mixed to each multiplex and the volume was reduced to 10 μl by evaporation.

6.4 **High-pH pre-fractionation and Liquid chromatography coupled to tandem mass spectrometry (nanoLC-MS/MS)**

For the analysis of secretome, high-pH (pH = 10) pre-fraction of TMT multiplexes was performed using High pH Reversed-Phase Peptide Fractionation kit (Thermo Fisher Scientific) and all collected fractions were evaporated to dryness. Collected fractions were redissolved in 22.5 μl of 0.1% trifluoroacetic acid (TFA), 2% AcN and 1 μl (~ 1 000 ng) was injected on an UltiMate 3000 RSLCnano System (Thermo Fisher Scientific) in two technical replicates. The analytical system consisted of a PepMap100 C18, 3 µm, 100 Å, 75 µm × 20 mm trap column and a PepMap RSLC C18, 2 µm, 100 Å, 75 µm × 250 mm analytical column (both from Thermo Fisher Scientific). The samples were loaded onto the trap column at a flow rate of 5 µl/min of 0.1% TFA, 2% AcN for 5 min. Tryptic peptides were separated via a linear gradient running from 2% to 45% of 0.1% formic acid, 80% AcN at a flow rate 250 nl/min for 80 min. Eluted peptides were sprayed into a Q Exactive Plus mass spectrometer using a Nanospray Flex ion source (Thermo Fisher Scientific) at 1.8 kV spray voltage for 107 min. Positive ion full scan MS spectra were acquired in the range of m/z 350-1 600, with 3×106 AGC target in the orbitrap at a resolution of 70 000 with a maximum ion injection time of 100 ms. A lock mass of m/z 445.12003 ([C2H6SiO]6) was used for internal calibration of the mass spectra. The fragmentation (MS/MS) spectra were acquired for the 10 most intense precursors. Isolation window of 1.6 m/z and normalized collision energy of 33 was used. Each fragmentation spectrum was acquired at a resolution of 35 000, with a 1×105 AGC target and a maximum ion injection time of 60 ms. The first mass was fixed to 100 m/z.

6.5 **MS Data acquisition and evaluation**

Recorded MS and MS/MS spectra were processed and searched in Proteome Discoverer 3.0 against a reviewed UniProt human reference protein database. Trypsin digestion specificity with up to two missed cleavages was used. Cysteine thiomethylation was set as a fixed modification and oxidation of methionine and proline was selected as a variable modification. The mass tolerance in MS and MS/MS mode was left at the default value for the initial search and 6 ppm was set as mass tolerance in MS mode for the main search. The false discovery rate for protein identification was left at the default value. Output files from Proteome Discoverer were processed in Perseus and R statistical environment v. 4.4.1 [4]. Three samples from collected conditioned E6 media (n=146) were excluded because of peptide loss during the sample preparation procedure. All remaining samples included in TMT protein quantification were randomized over TMT multiplexing. Potential technical deviation in individual TMT channels were removed using sum normalization of reporter ion intensities for each TMT multiplex (Supplementary Methods Figure 1 A-D) [24]. Technical variation among TMT multiplexes was removed using normalization to local pooled standard. Reporter ion intensities were firstly log2-transformed and subtracted from log2-transformed reporter intensity of proteins in the appropriate pooled reference standard (Supplementary Figure 1 E, F). The results of TMT normalization were illustrated using factoextra [25] FactoMineR [26], and ggplot2 [10] packages in R statistical environment v. 4.4.1 [4]. Subsequently, the entire dataset was divided into groups according to age and treatment of COs that theE6 media was harvested from as follows: 1) WT and PSEN1/2 mutant COs (D60), HSV-1 infection, 2) WT and PSEN1/2 mutant COs (D100), HSV-1 infection, 3) WT and PSEN1/2 mutant COs (D60), TBEV infection 4) WT and PSEN1/2 mutant COs (D100), TBEV infection. Control WT and PSEN1/2 mutant COs media (D60, D100) were added to each treatment group and designed datasets were filtered for valid values to exclude batches with less than 2 valid values (n=3-4). Differential expression LIMMA analysis [27] was performed in R statistical environment with Benjamin-Hochberg false discovery rate (FDR) correction at the significance level of 0.05. In each treatment, secreted protein levels in four groups were compared (WT-NTR, WT-inf, PSEN1/2 mutant-NTR, PSEN1/2 mutant-inf). All quantified proteins were classified according to their presence or absence in SASP atlas [28] significant difference in individual PSEN1/2 mutant/WT groups (B-H FDR p-value < 0.05), and down/upregulation based on log2FC value. For further analyses differentially secreted proteins (DSPs) with p.adj<0.1, |log2FC|>0.5 were used; they were assigned an ENTREZ ID further used in enrichment analyses as described in the 5.1 **Supplementary Methods** section, for DSPs with multiple mappings the mean_log2FC and combined_p.adj (Fisher’s method) were counted.

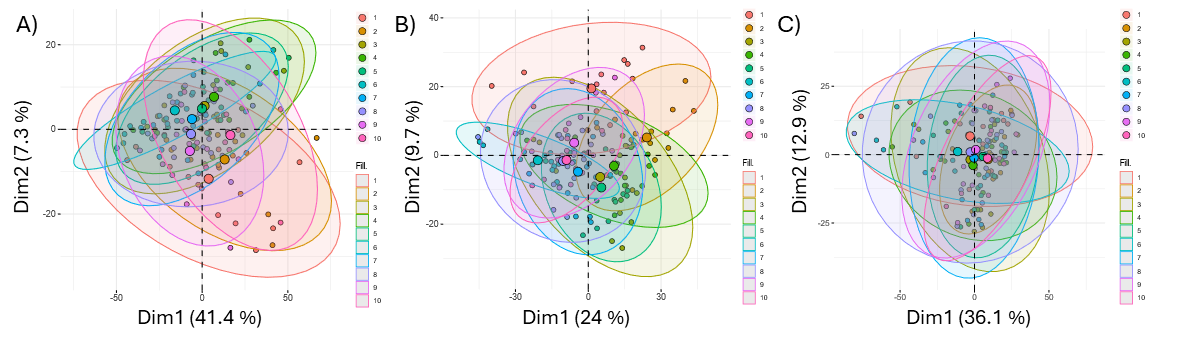

**Supplementary Figure 1: A, B)** Removal of technical variability of individual TMT channels by sum normalization of reporter ion intensities before normalization **(A)** and after normalization **(B)**. Removal of technical variability over the TMT multiplex by normalization to pooled reference standard **(C)**.

7. **References**

[10]	Wickham H. ggplot2: elegant graphics for data analysis. Second edition. Cham: Springer international publishing; 2016.
