## Supplementary_Tables_A-D for "Viral Infection Induces Alzheimer’s Disease-Related Pathways and Senescence in iPSC-Derived Neuronal Models"

**Table A: Cell lines used**

| **cell line name in the manuscript** | **cell line identifier (alternative identifier)** | **genotype** | **publication about derivation and characterization** |
| --- | --- | --- | --- |
| I3N iPSCs | I3N iPSCs | Wild-type (apparently healthy control) | Fernandopulle et al., 2018 |
| WT1 | MUNIi008-A (fWT1) | Wild-type (apparently healthy control) | Raska et al., 2021 |
| WT2 | MUNIi009-A (fWT2) | Wild-type (apparently healthy control) | Raska et al., 2021 |
| WT3 | MUNIi010-A (fWT3) | Wild-type (apparently healthy control) | Raska et al., 2021 |
| PSEN1/2 mutant 1 | MUNIi005-A (fAD1) | PSEN1 A243E | Raska et al., 2021 |
| PSEN1/2 mutant 2 | MUNIi006-A (fAD2) | PSEN1 A246E | Raska et al., 2021 |
| PSEN1/2 mutant 3 | MUNIi007-A (fAD3) | PSEN2 I144N | Raska et al., 2021 |

**Table B: Number of replicates per experiment**

| **figure** | **experimental method** | **used cell line, condition** | **total No. of biological replicates from all cell lines (n)** | **total No. of individual samples** | **sample type** | **number of samples per timepoint (shown in the figure)** | | |
| --- | --- | --- | --- | --- | --- | --- | --- | --- |
|  |  |  |  |  |  | **D21/D24** | **D60** | **D100** |
| 1D-F | IHC | i3N iPSCs_NTR | 6 | 18 | coverslips with 2D neurons | 18 |  |  |
|  |  | i3N iPSCs_HSV-1 | 3 | 3 | coverslips with 2D neurons | 3 |  |  |
|  |  | i3N iPSCs_TBEV | 4 | 8 | coverslips with 2D neurons | 8 |  |  |
| S1A | WB | i3N iPSCs_NTR | 4 | 4 | wells with 2D neurons | 4 |  |  |
|  |  | i3N iPSCs_HSV-1 | 4 | 4 | wells with 2D neurons | 4 |  |  |
| S1A | qRT-PCR | i3N iPSCs_NTR | 3 | 6 | wells with 2D neurons | 6 |  |  |
|  |  | i3N iPSCs_TBEV | 3 | 6 | wells with 2D neurons | 6 |  |  |
| S1B | IHC | i3N iPSCs_NTR | 6 | 18 | coverslips with 2D neurons | 18 |  |  |
|  |  | i3N iPSCs_HSV-1 | 3 | 3 | coverslips with 2D neurons | 3 |  |  |
|  |  | i3N iPSCs_TBEV, multiple MOIs | 4 | 25 | coverslips with 2D neurons | 5-8 |  |  |
| 2C-E | IHC | WT1, WT2, WT3_NTR | 3 | 65 | organoid sections |  | 31 | 34 |
|  |  | WT1, WT2, WT3_HSV-1 | 3 | 53 | organoid sections |  | 26 | 27 |
|  |  | WT1, WT2, WT3_TBEV | 3 | 54 | organoid sections |  | 30 | 24 |
| S2 | qRT-PCR | WT1, WT2, WT3_NTR | 3 | 12 | single organoids |  | 12 |  |
|  |  | WT1, WT2, WT3_HSV-1 | 4 | 16 | single organoids |  | 16 |  |
|  |  | WT1, WT2, WT3_TBEV | 3 | 12 | single organoids |  | 12 |  |
|  |  | PSEN1/2 mutant 1, 2, 3_NTR | 3 | 12 | single organoids |  | 12 |  |
|  |  | PSEN1/2 mutant 1, 2, 3_HSV-1 | 3 | 12 | single organoids |  | 12 |  |
|  |  | PSEN1/2 mutant 1, 2, 3_TBEV | 3 | 12 | single organoids |  | 12 |  |
| 3C-E | IHC | WT1, WT2, WT3_NTR | 3 | 31 | organoid sections |  | 31 |  |
|  |  | WT1, WT2, WT3_NTR_Aβ40 | 3 | 22 | organoid sections |  | 22 |  |
|  |  | WT1, WT2, WT3_NTR_Aβ42 | 3 | 18 | organoid sections |  | 18 |  |
|  |  | WT1, WT2, WT3_HSV-1 | 3 | 26 | organoid sections |  | 26 |  |
|  |  | WT1, WT2, WT3_HSV-1_Aβ40 | 3 | 22 | organoid sections |  | 22 |  |
|  |  | WT1, WT2, WT3_HSV-1_Aβ42 | 3 | 18 | organoid sections |  | 18 |  |
|  |  | WT1, WT2, WT3_TBEV | 3 | 30 | organoid sections |  | 30 |  |
|  |  | WT1, WT2, WT3_TBEV_Aβ40 | 3 | 22 | organoid sections |  | 22 |  |
|  |  | WT1, WT2, WT3_TBEV_Aβ42 | 3 | 18 | organoid sections |  | 18 |  |
| S3 | ELISA | i3N iPSCs | 5 | 5 | medium from wells of 2D neurons | 5 |  |  |
|  |  | WT1, WT2, WT3 | 3 | 16 | medium from single organoids | 16 | 8 | 8 |
| 4A-D, S4, 6E, S6B-D, 7B | 3'mRNA-seq | WT1, WT2, WT3_NTR | 3 | 12 | single organoids |  | 3 pools |  |
|  |  | WT1, WT2, WT3_HSV-1 | 3 | 12 | single organoids |  | 3 pools |  |
|  |  | WT1, WT2, WT3_TBEV | 3 | 12 | single organoids |  | 3 pools |  |
|  |  | PSEN1/2 mutant 1, 2, 3_NTR | 3 | 12 | single organoids |  | 3 pools |  |
|  |  | PSEN1/2 mutant 1, 2, 3_HSV-1 | 3 | 12 | single organoids |  | 3 pools |  |
|  |  | PSEN1/2 mutant 1, 2, 3_TBEV | 3 | 12 | single organoids |  | 3 pools |  |
| 5A-G, S5A-B, 6G-I, S6E | MS | WT1, WT2, WT3_NTR | 4 | 26 | single organoids |  | 12 | 14 |
|  |  | WT1, WT2, WT3_HSV-1 | 3 | 24 | single organoids |  | 12 | 12 |
|  |  | WT1, WT2, WT3_TBEV | 3 | 24 | single organoids |  | 12 | 12 |
|  |  | PSEN1/2 mutant 1, 2, 3_NTR | 3 | 24 | single organoids |  | 12 | 12 |
|  |  | PSEN1/2 mutant 1, 2, 3_HSV-1 | 3 | 24 | single organoids |  | 12 | 12 |
|  |  | PSEN1/2 mutant 1, 2, 3_TBEV | 3 | 24 | single organoids |  | 12 | 12 |
| 6B-C, S6A | IHC | PSEN1/2 mutant 1, 2, 3_NTR | 3 | 58 | organoid sections |  | 31 | 27 |
|  |  | PSEN1/2 mutant 1, 2, 3_HSV-1 | 3 | 57 | organoid sections |  | 30 | 27 |
|  |  | PSEN1/2 mutant 1, 2, 3_TBEV | 3 | 58 | organoid sections |  | 31 | 27 |

**Table C: Antibodies and reaction conditions for Western blotting and IHC**

| **antibody** | **cat. number** | **manufacturer** | **method** | **dilution** |
| --- | --- | --- | --- | --- |
| DAPI | D9542 | Merck | IHC | 1:1000/blocking buffer |
| Hoechst | H1399 | Thermo Fisher Scientific | IHC | 1:1000/blocking buffer |
| TUJ | 5568S | Cell Signaling Technology | IHC | 1:200/blocking buffer |
| MAP2 | AB5543 | Merck | IHC | 1:500/blocking buffer |
| β-Actin (8H10D10) | 3700 | Cell Signaling Technology | WB | 1:500/5 % milk |
| Anti-HSV 1 ICP4 | Ab6514 | Abcam | WB, IHC | 1:500/5 % milk |
| β-Amyloid (B-4) | sc-28365 | Santa Cruz Biotechnology | WB | 1:500/5 % milk |
| β-Amyloid (D54D2) | 8243S | Cell Signaling Technology | IHC | 1:200/blocking buffer |
| stably expressed mCherry of the TBEV construct | mCherry  autofluorescence | Haviernik et al., 2021 | IHC | none |
| Donkey Anti-Chicken AF647 | 703-606-155 | Jackson ImmunoResearch | IHC | 1:400/blocking buffer |
| Donkey Anti-Mouse AF568 | A10037 | Invitrogen | IHC | 1:400/blocking buffer |
| Donkey Anti-Rabbit AF488 | A21206 | Invitrogen | IHC | 1:400/blocking buffer |
| Donkey Anti-Rabbit AF568 | A10042 | Invitrogen | IHC | 1:400/blocking buffer |
| Donkey Anti-Rabbit AF647 | A31573 | Invitrogen | IHC | 1:400/blocking buffer |

**Table D: Primers for qRT-PCR**

| **primer** | **primer sequence** |
| --- | --- |
| HSV1-TK - Forward | GAAACTCCCGCACCTCTTCGG |
| HSV1-TK - Reverse | GGTTCCTTCCGGTATTGTCTCC |
| TBE virus gRNA/ TBEV-Forward | GGGCGGTTCTTGTTCTCC |
| TBE virus gRNA/ TBEV-Reverse | ACACATCACCTCCTTGTCAGACT |
